# Photoperiodic effects on early plant development in everbearing and seasonal flowering strawberry

**DOI:** 10.1101/2025.07.08.662500

**Authors:** Stephan David, Gabriela Ficová, Leo F.M. Marcelis, Julian C. Verdonk

## Abstract

In recent years, interest in vertical farming systems for strawberry cultivation has increased. Because of the increased costs associated with these systems, optimal use of resources, such as cultivation space and light, is a requirement. This necessitates a detailed understanding of plant development in different light conditions. The aim of this experiment was to analyze the effect of photoperiod on the early development of seasonal flowering and everbearing strawberry plants, and to compare the expression of genes known or suspected to be involved in the regulation of plant development between the two flowering types. Young, runner-propagated seasonal flowering and everbearing strawberry plants were grown under 10, 12, 14, or 16 h photoperiod in a climate-controlled vertical farm. Runners and flowers were removed throughout the experiment to limit their effects on assimilate balance. Morphological and physiological measurements were made weekly. Leaf material was collected for gene expression analysis. There were not many differences in plant growth and development between the short day conditions (10 and 12 h photoperiods), or long day conditions (14 and 16 h photoperiods). Short day conditions had a repressive effect on vegetative growth in all cultivars, but this effect was reduced in everbearing cultivars compared to seasonal flowering cultivars. No notable differences were found in gene expression between the two flowering types.

## INTRODUCTION

Strawberry (*Fragaria* x *ananassa* Duch.) is a popular and economically important fruit crop. The Netherlands produces around 80 million kg of strawberries yearly, in open soil, in plastic tunnels (polytunnels), and in greenhouses (Statistiek 2024). Strawberry yield has doubled since the year 2000. However, because more cultivation is done in greenhouses, which is more space efficient than open field cultivation, the area dedicated to strawberry production has not increased in the last decades (CBS 2024).

The relatively small size of the plant and the high market value and demand for the fruit makes strawberry a suitable candidate for vertical farm cultivation (Kouloumprouka Zacharaki et al. 2024). Vertical farms are indoor cultivation systems with controlled environments. Instead of sunlight, artificial is applied to drive photosynthesis. Typically, they contain vertically stacked growing layers or multiple cultivation floors (van Delden et al. 2021). These systems offer several benefits over cultivation in open field or greenhouses, such as year-round cultivation, independent of weather or climate, and a reduction in environmental pollution by pesticides and fertilizers (van Delden et al. 2021). However, the technology required for vertical farming systems cultivation result in high costs. This necessitates that, to be competitive, resources are optimally utilized, and yield is maximized. A detailed understanding of early plant development, and how it is influenced by different photoperiods, would facilitate this.

Strawberry cultivars can be distinguished based on flowering type. Seasonal flowering cultivars require exposure to short day conditions for flower initiation and produce a single flush of flowers (Heide 1977; Hytönen and Kurokura 2020). Exposure to long day conditions inhibits flower initiation but stimulates vegetative growth. In strawberry, short day and long day responses are determined by whether the photoperiod falls above or below a cultivar-specific critical day length (Heide et al. 2013; Stewart and Folta 2010). In commercial cultivation, after the fruit harvest, seasonal flowering plants are typically replaced by a later-flowering cultivar. Everbearing cultivars initiate flowers under long day conditions and as such do not require exposure to short day conditions for flower initiation (Rivero et al. 2021). This behavior is generally attributed to the dominant PERPETUAL FLOWERING (PF) locus, which abolishes photoperiod-dependent inhibition of flowering (Perrotte et al. 2016; Hytönen and Kurokura 2020). However, which locus is responsible for everbearing behavior is cultivar-dependent (Lewers 2019). They also produce fruits over an extended period compared to seasonal flowering types (De Camacaro et al. 2002).

The aim of this experiment is to analyze the effect of photoperiod on the early development of seasonal flowering and everbearing strawberry plants, and to compare the expression of genes known or suspected to be involved in developmental regulation between the two flowering types. Young runner propagated plants were grown in a vertical farming system under four different photoperiods (10, 12, 14, or 16 h). Plant morphology was analyzed weekly and after 10 weeks a destructive harvest was conducted. We analyzed the expression of *SOC1*, a transcription factor that promotes vegetative development and represses flowering through induction of the anti-florigen *TFL1* (Mouhu et al. 2013; Muñoz-Avila et al. 2022); *CO*, a key regulator of photoperiodic flowering in Arabidopsis and likely involved in strawberry photoperiodism (Kurokura et al. 2017; Muñoz-Avila et al. 2022); and *DAM3* and *DAM4*, candidate regulators of seasonal dimorphism in strawberry that respond to short day conditions (David et al. 2025).

## MATERIALS AND METHODS

### Plant material

Two seasonal flowering and two everbearing cultivars were used (Table 1). Plants were provided by breeding company Fresh Forward (Huissen, The Netherlands), where they were propagated from runners on June 1, 2023, and grown in plugs (125 ml coconut fiber, peat, and perlite; 40:40:20 ratio) under long day conditions in the greenhouse. On June 16, 2023, the plants were transported to Wageningen University. Prior to transport, plug plants were treated with a Controlled Atmosphere Temperature Treatment (CATT) as a standard integrated pest management measure (Verschoor et al. 2015). All leaves besides the two or three youngest were removed to standardize leaf area index and carbon assimilation capacity at the start of the experiment, as plants were received with varying numbers of leaves. Retaining this variation could have introduced growth differences unrelated to the photoperiod treatment. (Figure 1).

**Table 1.** the four cultivars which were used in the experiment. The column ‘name’ indicates how each cultivar is referred to in the current paper. Plants of each cultivar were propagated from runners and rooted in plugs.

| Cultivar | Flowering type | Name |
| --- | --- | --- |
| Elsanta | Seasonal | SF_Els |
| Fandango | Seasonal | SF_Fan |
| Favori | Everbearing | EB_Fav |
| D080-037-2020 | Everbearing | EB_D08 |

**Figure 1.**
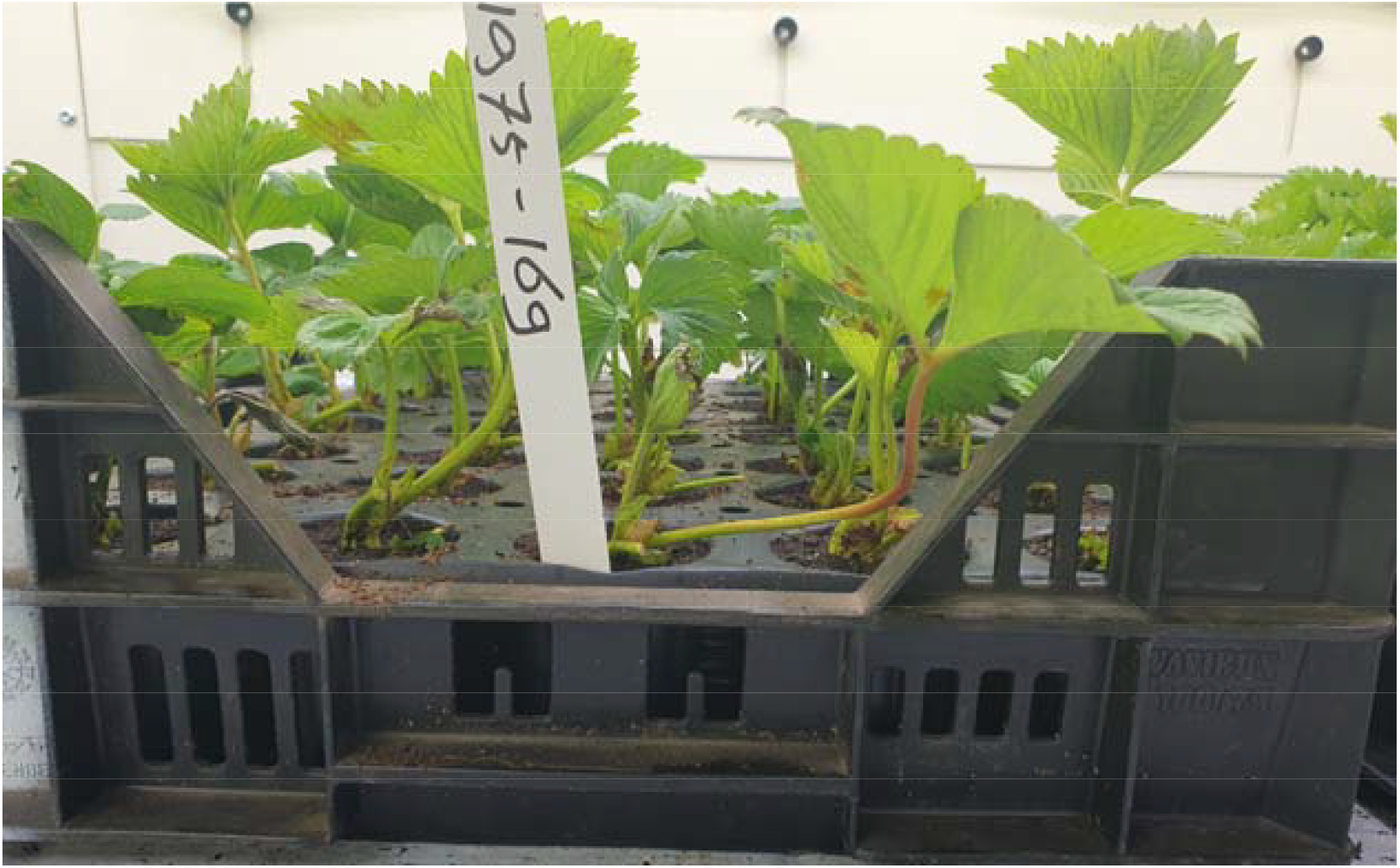
plug plants in a tray, after leaf removal. Plants were grown in plugs, see picture, under a 13-hour photoperiod for ten days in order to acclimate to the climate cell conditions. Picture taken on the day the plants were transported to the climate cell, on June 16, 2023.

### Growing conditions

The experiment was conducted in a climate cell (Wageningen University and Research). The climate cell contained three layers with four cultivation compartments on each layer, for a total of 12 compartments. The three cultivation layers served as independent blocks in the split-plot design, providing three true statistical replicates. Photoperiod treatments were assigned to compartments within each layer, with each of the four treatments represented once per layer (Fig 2B). Each compartment, measuring 250 by 80 cm, contained 16 plants, four of each cultivar. Measurements of four plants were averaged into one replicate, for a total of three replicates per treatment-cultivar combination (n=3). Light was provided with LED lamps (GPL PM 168 DRBWFR_R L120 3.1 C4, Signify, Eindhoven, the Netherlands). The spectral composition (400-499 nm: 23%; 500-599 nm: 37%; 600-699 nm: 27%; 700-799 nm: 13%) was measured with a spectroradiometer (model SS-110, Apogee Instruments, Logan, UT, USA) (Supplement 1). Plants were irrigated as needed using a standard strawberry nutrient solution applied by an ebb and flow system, with frequency adjusted throughout the experiment based on observed liquid uptake rates. Relative humidity and temperature in each compartment were continuously measured with a sensor (Onset, HOBO MX2300) (Supplements 2-5). Relative humidity was initially set to 75% day and night. On July 10, 2023, settings were adjusted to 60% during the day and 90% at night, with a peak of 100% at 08:00, to manage tip burn. The plants were acclimated to the conditions in the climate cell by growing them under a 13 h photoperiod (231 µmol·m^-2^·sec^-1^) for ten days before the treatment period. On June 23, 2023, from each cultivar, 48 plants were selected resulting in a total of 192 plants across the experiment. Plants were potted in 2 L pots with coconut fiber, peat, and perlite (40:40:20 ratio) (Figure 2).

**Figure 2.**
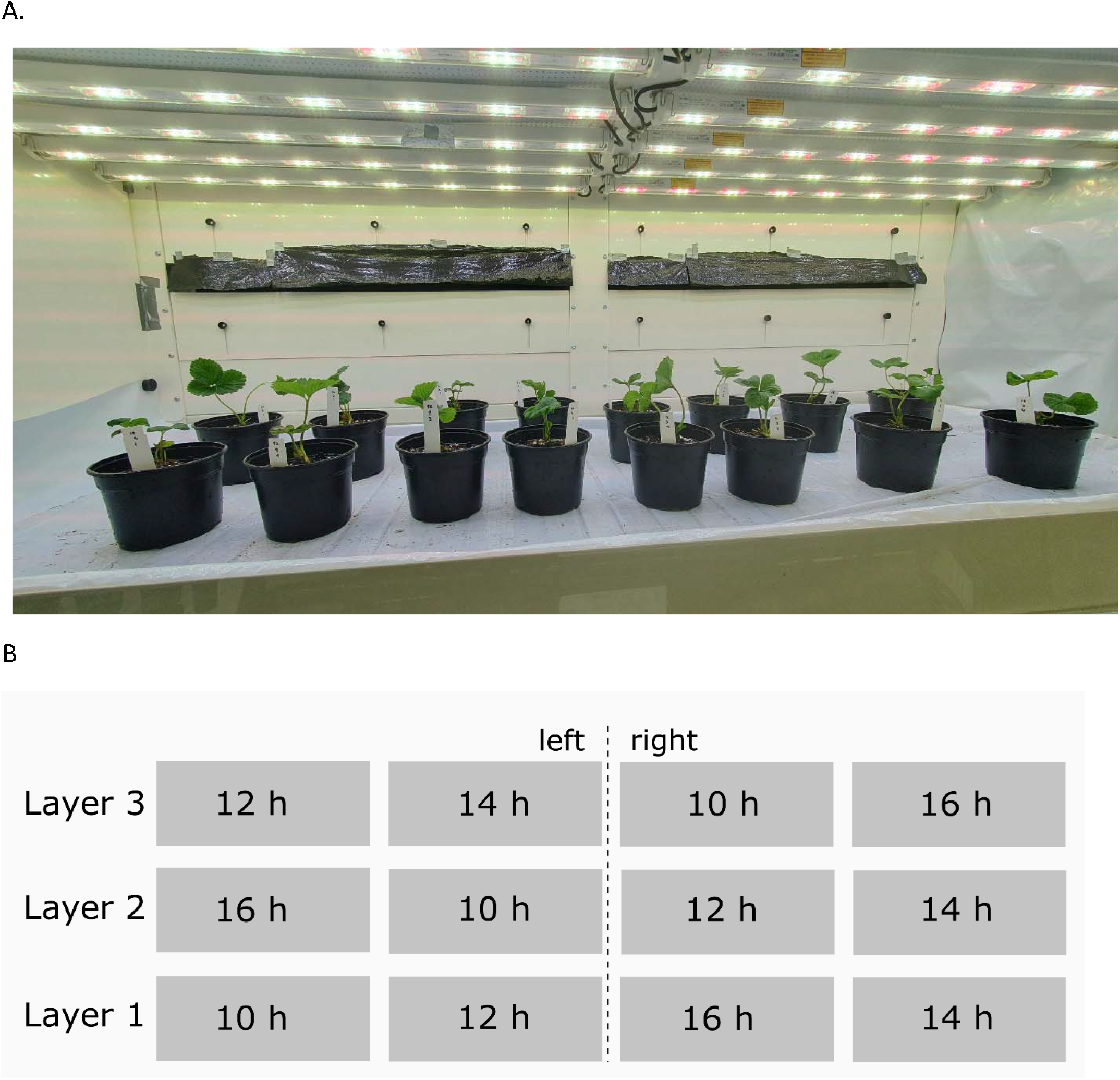
A, one of the climate cell compartments after potting and placement of the plug plants. Potted plants were divided over 12 compartments before the start of the treatment period. June 23, 2023. B, schematic overview of the climate cell layout. The climate cell consisted of four compartments per growing layer, for a total of twelve compartments. The four photoperiod treatments were randomly distributed over each growing layer, which served as blocks in the split-plot design. Each compartment contained four plants of each of the four cultivars during the experimental period.

### Treatments

On June 26, 2023, photoperiod treatments were started. There were four different treatments consisting of a 10, 12, 14 or 16 h photoperiod. The light intensity was adjusted so that each treatment had a daily light integral (DLI) of 10.8 mol·m^-2^·d^-1^ (Table 2). The treatment period lasted until September 13, 2023.

**Table 2.** overview of the treatments and light intensities. The DLI was the same for all treatments (10.8^−2^ mol.m^−1^

| Photoperiod | Light intensity<br>($\mu\text{mol} \cdot \text{m}^{-2} \cdot \text{sec}^{-1}$) | DLI<br>($\text{mol} \cdot \text{m}^{-2} \cdot \text{day}^{-1}$) |
| --- | --- | --- |
| 10 | 300 | 10.8 |
| 12 | 250 | 10.8 |
| 14 | 214 | 10.8 |
| 16 | 188 | 10.8 |

### Measurements

At the start of the treatment period, five surplus plants of each cultivar were selected for (destructive) analysis of the crowns and meristems, also known as flower mapping. In this process, the number of crowns, as well as the location and developmental stage of the meristem positions were described. These measurements were performed at Planta Logica (Ede, the Netherlands). During the treatment period of the experiment, vegetative development and flowering were analyzed. On a weekly basis, the youngest fully developed leaf was selected (defined as the second youngest, fully opened leaf), of which the following measurements were taken: The width of the left leaf lobe and the length of the middle lobe; The length of the petiole; Chlorophyll content was estimated with an optical sensor (Force-A, Dualex, Orsay, France). Leaf area (LA) was calculated from individual leaf lobe measurements, according to the method described by Demirsoy *et al*. (Equation 1) (Demirsoy et al. 2005).

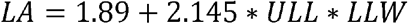

**Equation 1, estimation of leaf area from lobe measurements. LA is the leaf area of the whole leaf (cm). ULL is the upper lobe length (cm). LLW is the left lobe width (cm). Based on Demirsoy *et al*. (2005) (Demirsoy et al. 2005)**.

Additionally, the number of runners and open flowers was recorded. Runners were removed after counting. Flowers were removed as soon as or before they reached the small green stage of fruit development, to limit the effect on the assimilate balance. At the end of the experiment, on September 13, 2023, the final destructive measurements were done. Of each plant, the total number of leaves, total leaf area, fresh and dry weight (dried in the oven at 105°C for 48 hours) of all leaves was measured. Afterwards, four plants of each of the seasonal flowering cultivars were selected for flower mapping. Everbearing cultivars were not included in this analysis, as both cultivars already carried generative meristems at the start of the treatment period (Figure 9), and dissection was therefore not expected to yield additional information on photoperiod-induced flower initiation.

### Leaf sample collection

Leaf material was taken on June 27, July 18, August 8, and August 29, 2023 (weeks 1, 4, 7 and 10 of the treatment period). Two 1 cm diameter leaf disks were taken from two leaf lobes of the second youngest leaf. Leaf material of plants of the same cultivar in the same compartment was combined and subsequently flash frozen in liquid nitrogen. All leaf material was collected within the first hour after the lights switched on.

### RNA extraction

Leaf material frozen in liquid nitrogen was finely ground in a ball mill grinder (Retsch, MM400, Haan, Germany). RNA was extracted from the plant samples using the CTAB method (adapted from Schultz *et al*. 1994 (Schultz et al. 1994)) with lithium chloride precipitation. All centrifuge steps were done in a microcentrifuge at maximum speed. In a 2 mL tube, 750 µL extraction buffer (1.4M NaCl, 20 mM EDTA pH8, 100 mM TRIS pH8, 2% CTAB, 2% PVP-40, 1% β-mercaptoethanol, milli-Q water (MQ) to 750 µL) was added to ±50 mg of frozen leaf powder. Samples were incubated 15 min at 65°C. 750 µL chloroform was added and samples were centrifuged at max speed for 5 min. The supernatant was transferred to a tube with 500 µL isopropanol, homogenized and centrifuged at max speed for 5 min. Supernatant was removed and pellet was washed with 70% ethanol. The pellet was dissolved in 205 µL MQ and RNA was precipitated by adding 67 µL 8M LiCl and incubating for 30 min at −20°C. The RNA was pelleted by centrifugation at max speed for 30 min at 4°C. Pellet was washed once more with 70% ethanol, air dried for 5 minutes and dissolved in 50 µL MQ. RNA quantity and quality were assessed with spectrophotometric analysis and gel-electrophoresis.

### cDNA synthesis

cDNA was synthesized from RNA using the High-Capacity cDNA Reverse Transcription Kit (#4368813, Applied Biosystems, Waltham, MA, USA). In a PCR tube, 200 ng total RNA was combined with 2 µL 10x reaction buffer, 2 µL random primer mix, 0.8 µL 10 mM dNTP mix, 1 µL RTase enzyme, and MQ to 20 µL. Reverse-transcriptase PCR was done in a thermal cycler (SensoQuest GmbH, Labcycler Gradient, Göttingen, Germany) with the following program: 10 min at 25°C, 120 min at 37°C, 5 min at 85°C. Finally, cDNA was diluted by adding 80 µL MQ. Quality of the cDNA was assessed by PCR amplification of *ACTIN1* and gel-electrophoresis.

### qRT-PCR

For each primer pair (gene of interest or reference gene, Table 3) a primer mix was prepared (1 µL 10 µM forward primer, 1 µL µM reverse primer, 10 µL iQ SYBR Green Supermix (Bio-Rad, Hercules, CA, USA) per reaction). For each cDNA sample, a cDNA mix was prepared (1 µL cDNA, 7 µL MQ). qRT-PCR reactions were prepared by combining 12 µL primer mix with 8 µL cDNA mix in a 96-well plate (BIOPlastics, B17489, Landgraaf, the Netherlands). Plates were analyzed in the CFX Thermo Cycler (Bio-Rad), following a three-step program (initial denaturation for 3 min at 95°C, 40 cycles of 95°C for 10 sec, 30 sec at annealing temperature, and 30 seconds at 72°C, followed by a melt curve from 95 to 55°C). Raw data was analyzed with the LinRegPCR software (version 2021.2) and variation between plates was reduced with FactorqPCR (version 2020.0) (Ruijter et al. 2009, 2015).

**Table 3.**
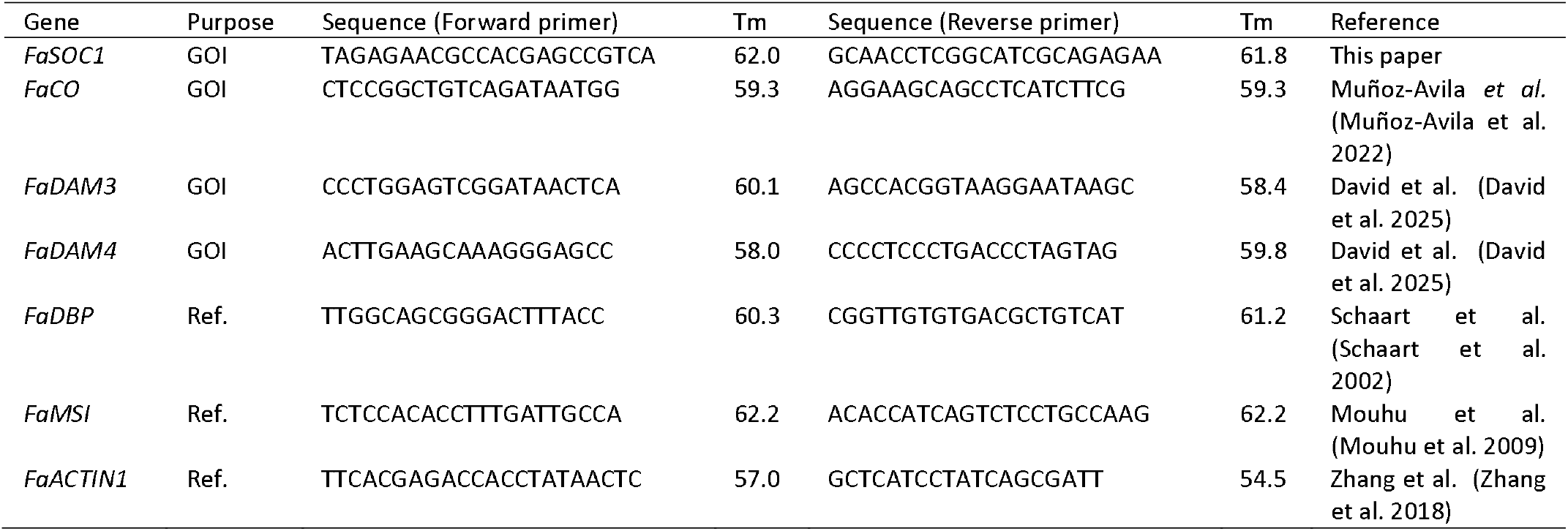
list of primers that were used for cDNA quality assessment with PCR and for expression analysis with qRT-PCR. GOI: gene of interest; Ref.: qRT-PCR reference gene. *FaSOC1*: *SUPPRESSOR OF OVEREXPRESSION OF CONSTANS 1*, repressor of flowering and promoter of vegetative growth; *FaCO*: *CONSTANS*, regulator of photoperiodic flowering; *FaDAM3* and *FaDAM4*: *DORMANCY-ASSOCIATED MADS-box* genes, candidate regulators of seasonal dimorphism; *FaDBP, FaMSI*, and *FaACTIN1*: reference genes used for expression normalization.

### Statistical analysis

The experiment was set up in a split-plot design with blocks, with photoperiod as the main plot and cultivar as the subplots. The growth chamber consisted of three cultivation layers, each of which was considered a block. Split-plot ANOVA was performed in Genstat (VSN International, v22.1.0.195). F-probabilities below 0.05 were considered significant. Fisher’s unprotected LSD test was applied for multiple comparisons. On the meristem analysis before the treatment period, a one-way ANOVA with cultivar as the factor was performed in SPSS Statistics (IBM, v28.0.1.1 (15)).

## RESULTS

We did not find any differences in any of the measured traits (petiole length, leaf area, and chlorophyll content) between the two short (10 and 12 h), or between the two long photoperiods (14 and 16 h) in any cultivar (Figure 3-5). Due to increasingly severe tip-burn from week 4 onwards in ‘SF_Fan’, the following section primarily focusses on the measurements of ‘SF_Els’, ‘EB_Fav’ and ‘EB_D08’.

**Figure 3.**
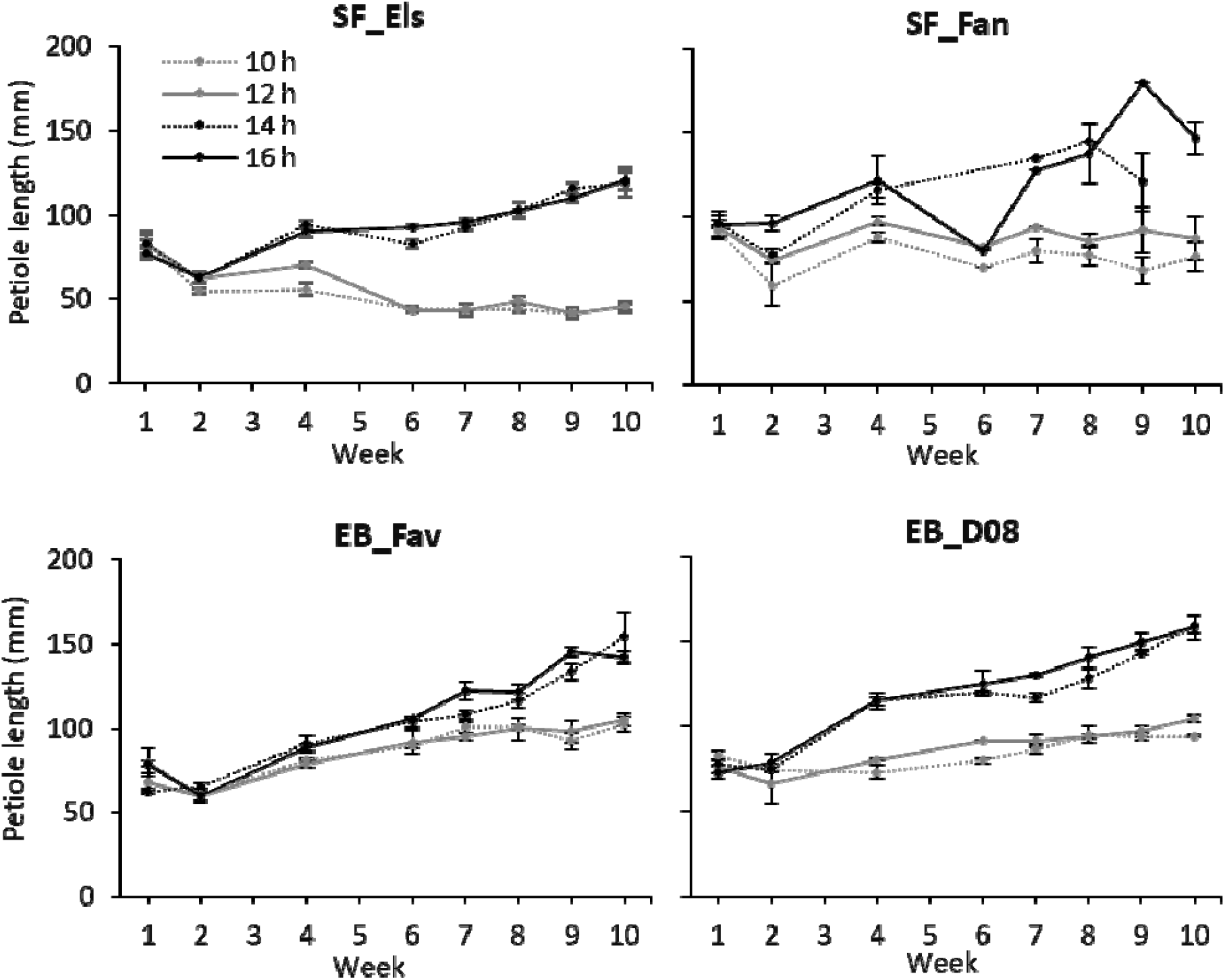
the effect of photoperiod on petiole length of newly emerged leaves of two seasonal flowering (SF) cultivars and two everbearing (EB) cultivars over time. SF_Els: Elsanta; SF_Fan: Fandango; EB_Fav: Favori; EB_D08: D080-037-2020. Each data point is the average of three plots, each plot consisting of four replicate plants per cultivar. Error bars denote the standard error. Photoperiods of 10 and 12 h represent short day (SD) conditions; 14 and 16 h represent long day (LD) conditions.

Day lengths of 14 and 16 h caused the petioles of newly emerged leaves to gradually lengthen over the treatment period in all cultivars (Figure 3). In ‘SF_Els’, the 10 and 12 h photoperiods resulted in a decrease in petiole length, especially between weeks 4 and 6. In the everbearing types (‘EB_Fav’ and ‘EB_D08’), petiole length did not decrease during the treatment period, although the petioles of plants in short day conditions were shorter compared to those of the plants in long day conditions.

In all cultivars, the long photoperiods increased the leaf area of newly emerged leaves compared to the short photoperiods (Figure 4). In ‘SF_Els’, the leaf size of plants in long photoperiods was considerably larger than in the everbearing types. The leaf size of plants in short photoperiods was similar between all cultivars, especially in the second half of the treatment period, after the effects of the pre-treatment had worn off.

**Figure 4.**
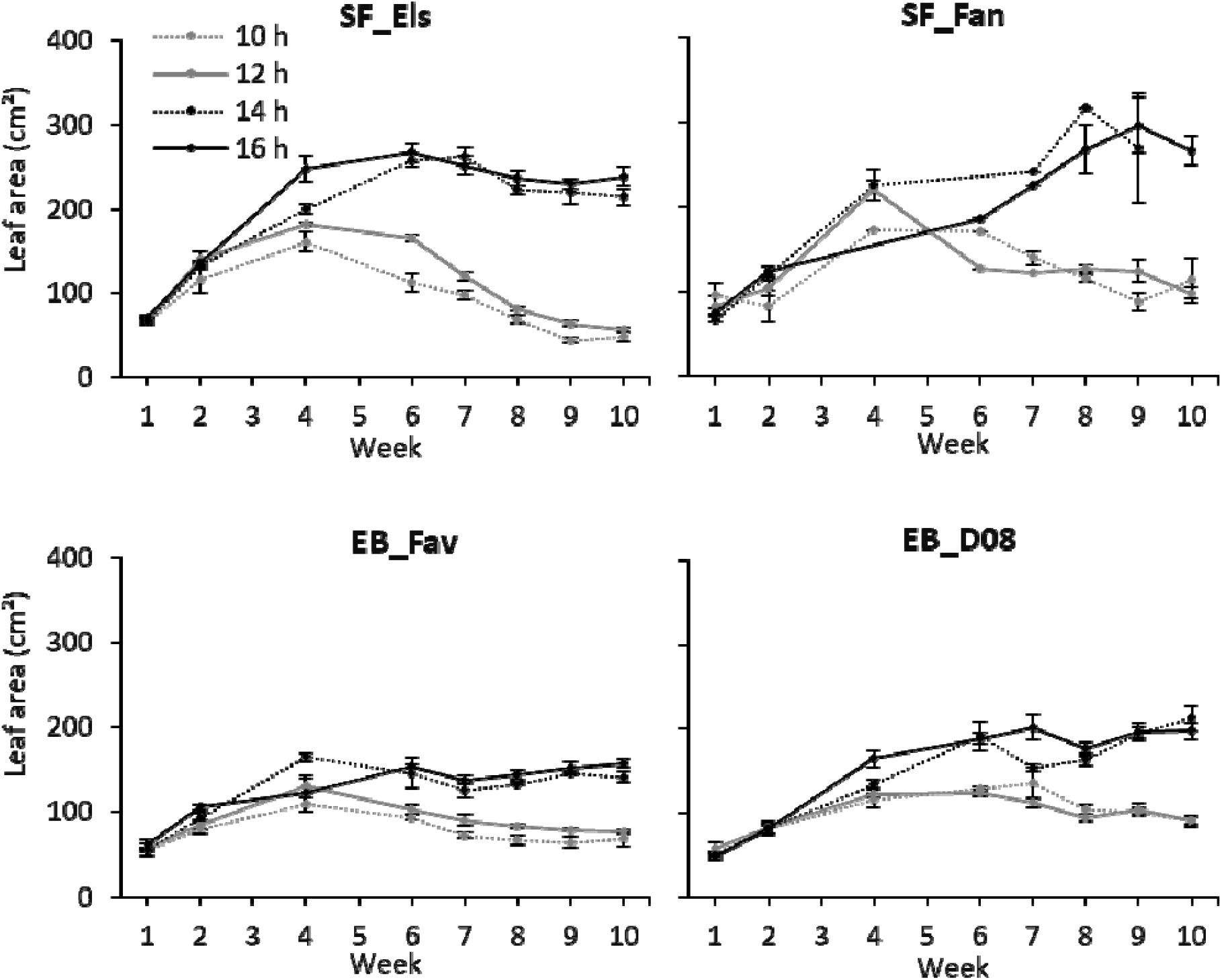
the effect of photoperiod on leaf area of newly emerged leaves of two seasonal flowering (SF) cultivars and two everbearing (EB) cultivars over time. SF_Els: Elsanta; SF_Fan: Fandango; EB_Fav: Favori; EB_D08: D080-037-2020. Each data point is the average of three plots, each plot consisting of four replicate plants per cultivar. Error bars denote the standard error. Photoperiods of 10 and 12 h represent short day (SD) conditions; 14 and 16 h represent long day (LD) conditions.

In all cultivars, short photoperiods increased leaf chlorophyll content compared to long photoperiods (Figure 5). Differences between long and short photoperiods became noticeable only in the second half of the treatment period.

**Figure 5.**
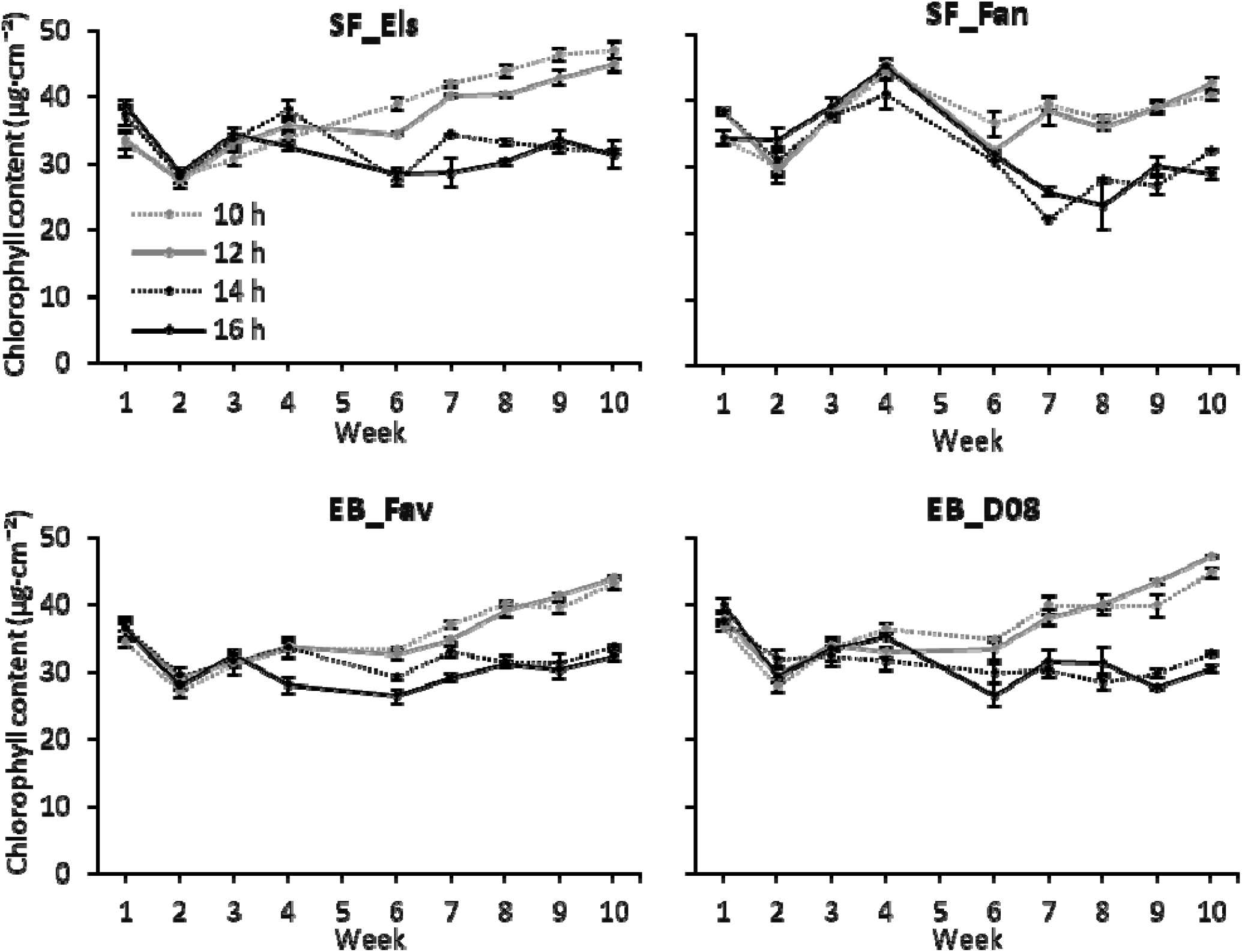
the effect of photoperiod on chlorophyll content of newly emerged leaves of two seasonal flowering (SF) cultivars and two everbearing (EB) cultivars over time. SF_Els: Elsanta; SF_Fan: Fandango; EB_Fav: Favori; EB_D08: D080-037-2020. Each data point is the average of three plots, each plot consisting of four replicate plants per cultivar. Error bars denote the standard error. Photoperiods of 10 and 12 h represent short day (SD) conditions; 14 and 16 h represent long day (LD) conditions.

All cultivars produced fewer runners under 10 and 12 h photoperiods compared to 14 and 16 h photoperiods (Figure 6). Under 10 and 12 h photoperiods, runner production in all cultivars was very low or completely halted before the end of the treatment period. Under 14 and 16 h photoperiods, some cultivars produced runners throughout the treatment period, such as ‘SF_Els’ under 16 h photoperiod, or ‘EB_D08’ under 14 h photoperiod. In ‘EB_Fav’, runner production peaked in week 5, after which it decreased until runner production had ceased by the end of the treatment period. A similar trend, peaking in week 5, could be seen in EB_D08, although less distinct.

**Figure 6.**
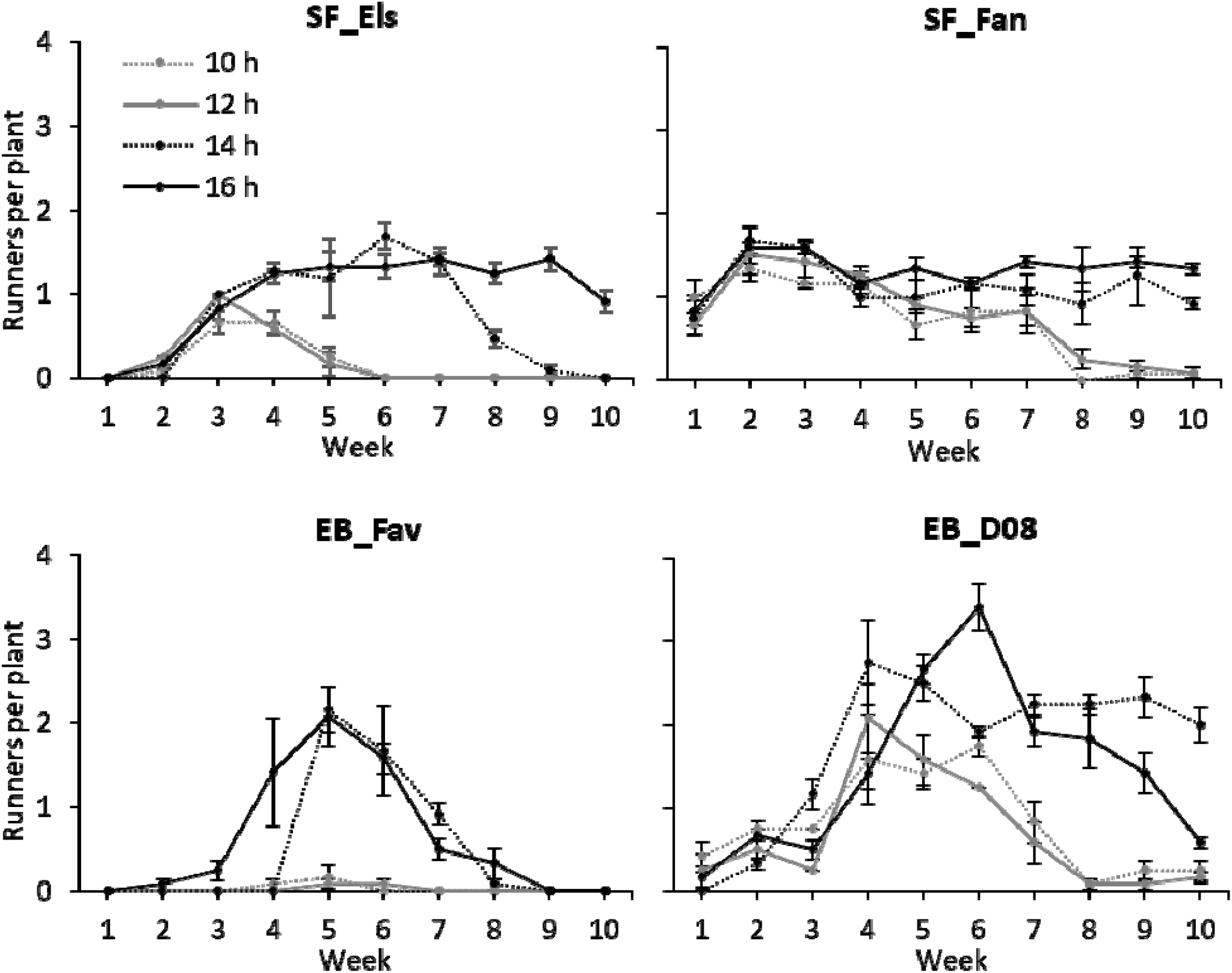
the effect of photoperiod on runner production of two seasonal flowering (SF) cultivars and two everbearing (EB) cultivars over time. SF_Els: Elsanta; SF_Fan: Fandango; EB_Fav: Favori; EB_D08: D080-037-2020. Each data point is the average of three plots, each plot consisting of four replicate plants per cultivar. Error bars denote the standard error. Photoperiods of 10 and 12 h represent short day (SD) conditions; 14 and 16 h represent long day (LD) conditions.

Plants of the cultivar SF_Els did not produce any flowers. ‘SF_Fan’ produced a few flowers under 12 h photoperiod in the last four weeks of the treatment period (Figure 7). Both everbearing cultivars flowered. Flowering started later in ‘EB_Fav’, but the rate of flower emergence was higher than in ‘EB_D08’. No photoperiodic effect was found on the starting time or rate of flower emergence in either everbearing cultivar.

**Figure 7.**
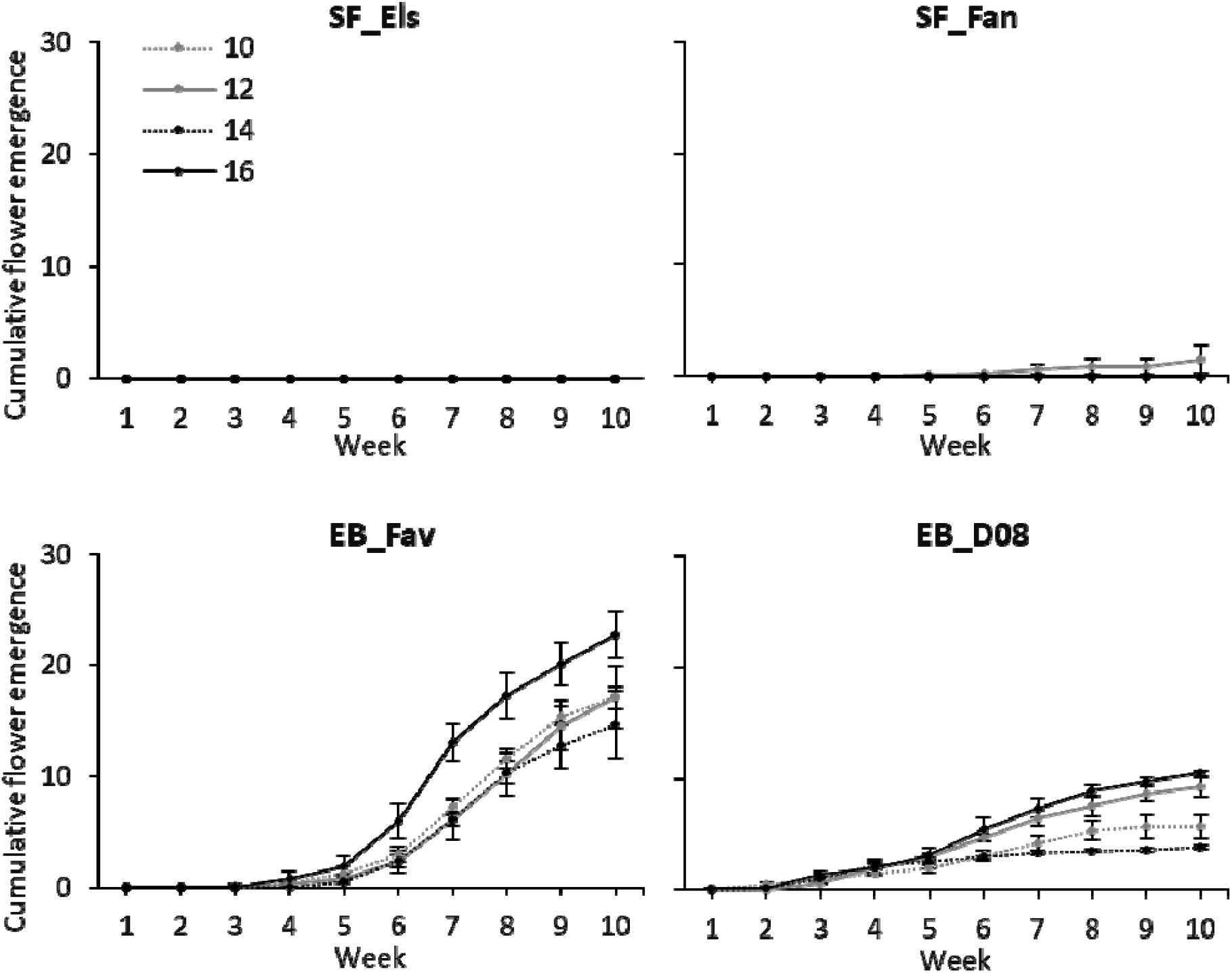
the effect of photoperiod on cumulative flower emergence of two seasonal flowering (SF) cultivars and two everbearing (EB) cultivars over time. SF_Els: Elsanta; SF_Fan: Fandango; EB_Fav: Favori; EB_D08: D080-037-2020. Each data point is the average of three plots, each plot consisting of four replicate plants per cultivar. Error bars denote the standard error. Photoperiods of 10 and 12 h represent short day (SD) conditions; 14 and 16 h represent long day (LD) conditions.

The destructive harvest data (Figure 8) provides the clearest overall picture of photoperiodic effects on vegetative biomass. It should be noted that runners and flowers were removed throughout the experiment, which limits sink competition but may have influenced absolute biomass values. In all cultivars, except ‘SF_Fan’, 14 and 16 h photoperiods resulted in an increase in total leaf area (Figure 8A). This increase is explained by an increase in leaf area per leaf under the same conditions (Figure 8B), which was observed in all cultivars, and not by an increase in number of leaves. In seasonal flowering cultivars the number of leaves decreased under 14 and 16 h photoperiods, compared to the 10 and 12 h photoperiods (Figure 8C). In the everbearing cultivars no such effect was found. Similar to total leaf area, the leaf dry weight increased in long photoperiods in all cultivars except ‘SF_Fan’ (Figure 8D). Photoperiod did not affect the leaf dry matter content or the specific leaf area of any cultivar (Figure 8E-F).

**Figure 8.**
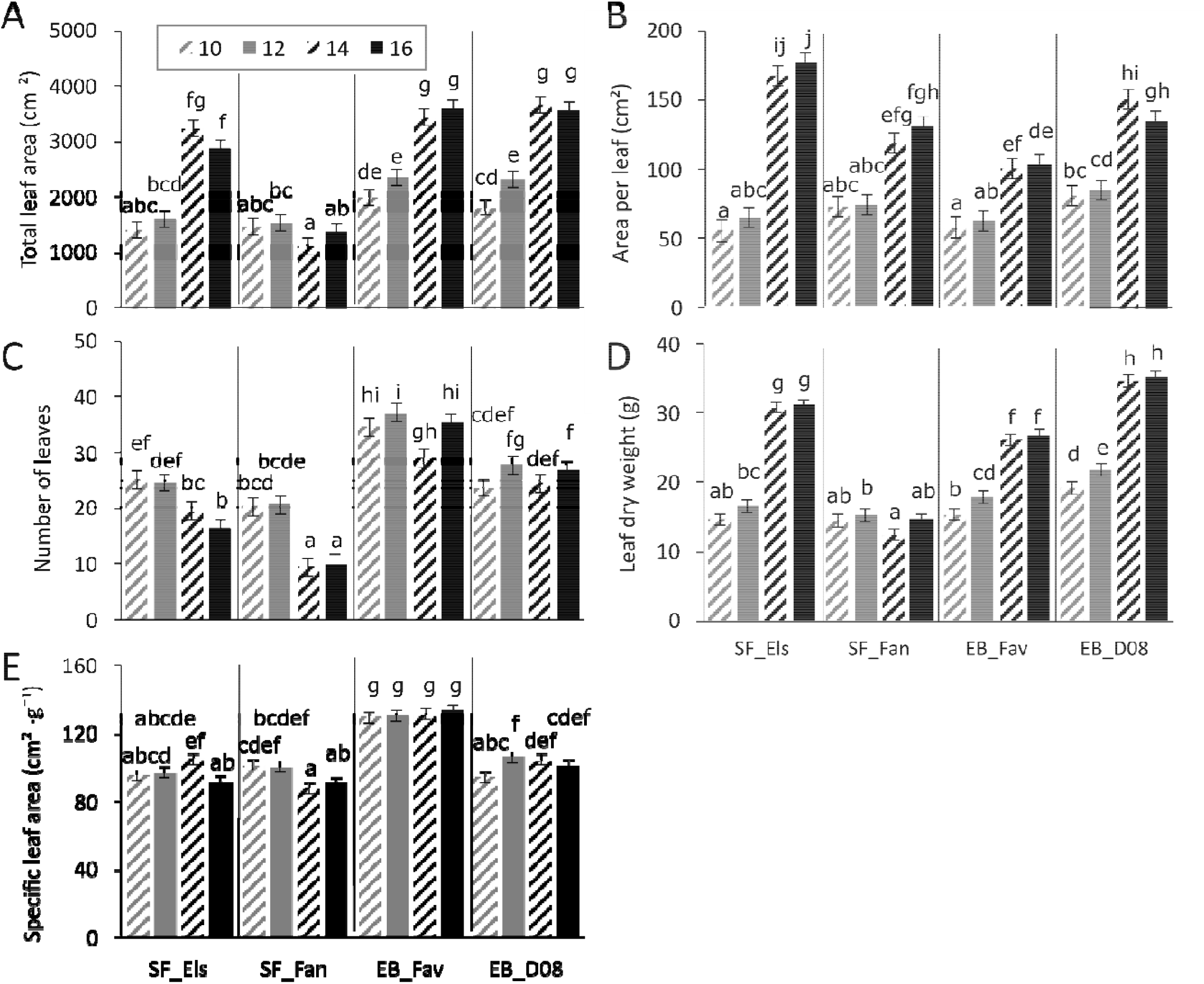
the effect of daylength on total leaf area (A), number of leaves (B), area per leaf (C), leaf dry weight (D), leaf dry matter content (E), and specific leaf area (F). After 10 weeks of photoperiodic treatment, vegetative growth was analyzed by destructive harvest. Total number of leaves, total leaf area, and leaf dry weight were measured. Aditionally, area per leaf, leaf dry matter content and specific leaf area were calculated. SF_Els: Elsanta; SF_Fan: Fandango; EB_Fav: Favori; EB_D08: D080-037-2020. Each data point is the average of three repetitions, each repetition consisting of four replicates. Data were analyzed with split-plot ANOVA.

Before the treatment period, all plants of each cultivar had one crown (Figure 9). The everbearing cultivars both contained generative meristems while the seasonal flowering cultivars were completely vegetative at this point. After the treatment period the seasonal flowering cultivars were analyzed again. Under 10 and 12 h photoperiod, the amount of crowns had increased (Figure 10A). In ‘SF_Els’, under 10 and 12 h photoperiod the number of positions was higher compared to long day 14 and 16 h photoperiod. In ‘SF_Fan’ no significant effect of photoperiod on number of meristem positions was found (Figure 10B). The average number of meristem positions per crown was lower under 10 and 12 h photoperiod (Figure 10C). The percentage of generative meristem was significantly higher under 10 and 12 h photoperiod in both cultivars, as the majority of plants did not develop generative meristem under 14 and 16 h photoperiod.

**Figure 9.**
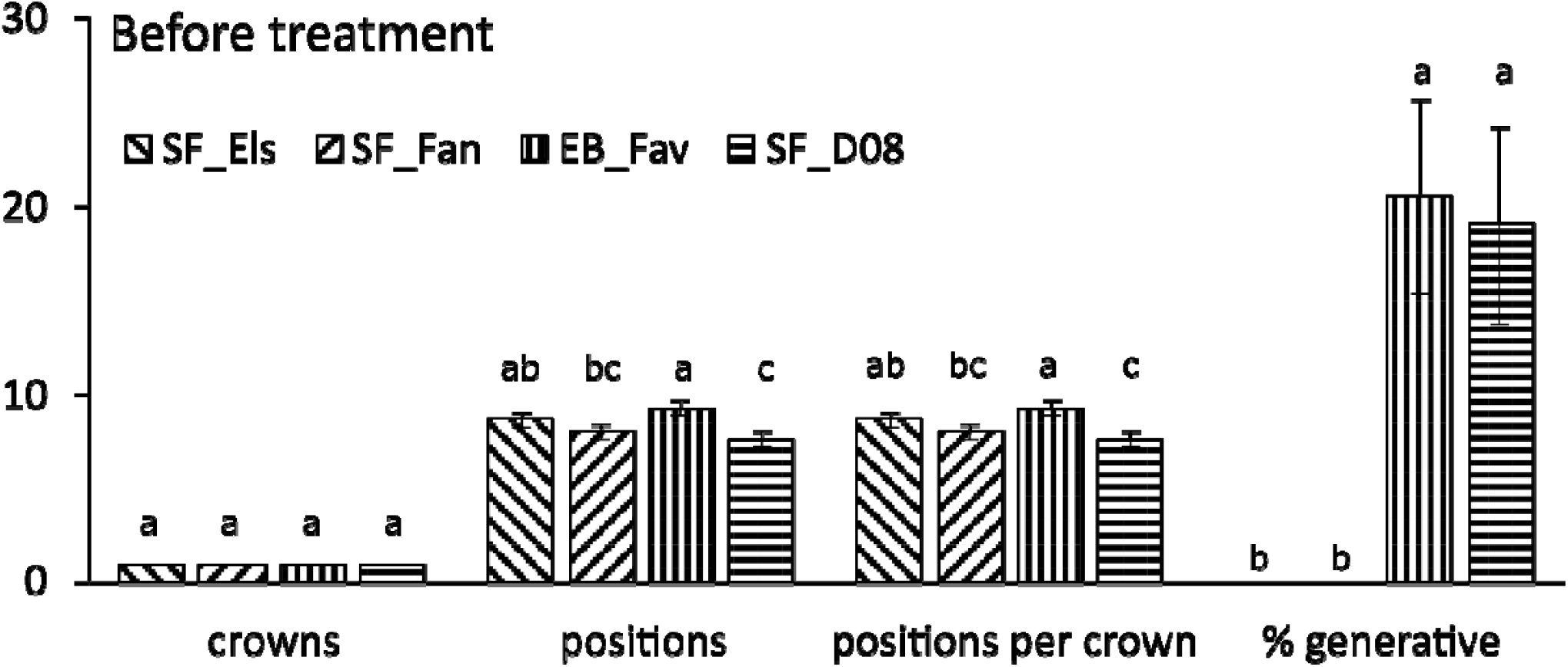
generative growth before the photoperiod treatment. The number of crowns per plant, number of meristem positions, and meristem status (vegetative or generative) were counted. SF_Els: Elsanta; SF_Fan: Fandango; EB_Fav: Favori; EB_D08: D080-037-2020. Each data point is the average of five repetitions, each repetition consisted of a single plant. Data were analyzed with one-way ANOVA. Different letters denote significant differences according to Fisher’s protected LSD.

**Figure 10.**
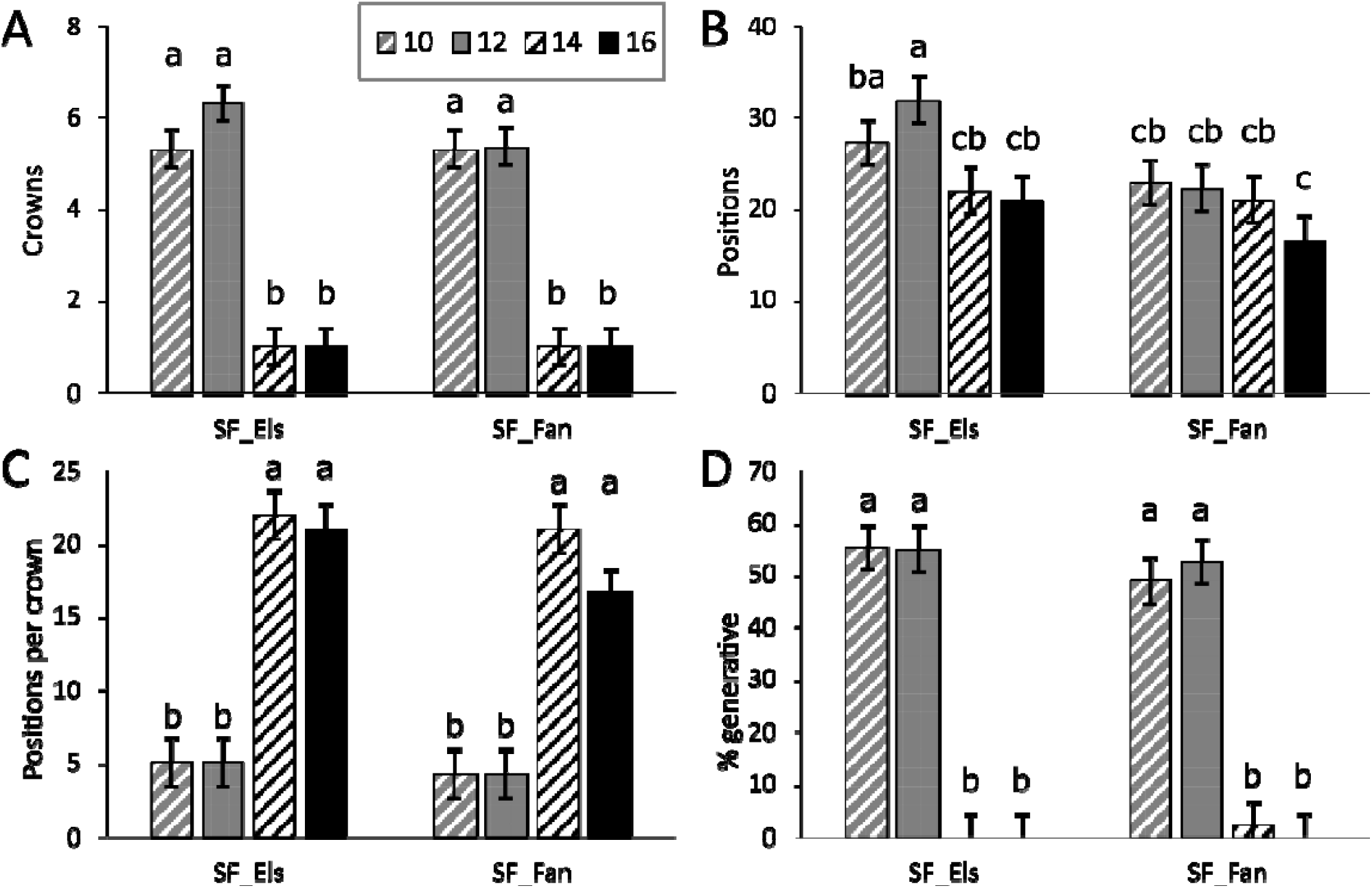
the effect of photoperiod on number of crowns per plant (A), number of meristem positions (B, C) and meristem status (vegetative or generative) (D). SF_Els: Elsanta; SF_Fan: Fandango. Each data point is the average of three repetitions, each repetition consisted of a single plant. Data were analyzed with split-plot ANOVA.

The expression of *SOC1* decreased under 10 h photoperiod in all cultivars, while under 16 h photoperiod *SOC1* expression remained relatively high. *CO* expression was found to vary over time and between cultivars and did not respond consistently to photoperiod, but generally *CO* expression was lower under the 16 h photoperiod. *DAM3* expression increased under 10 h photoperiod in the first seven weeks of the treatment period, after which it stabilized or decreased slightly. Under 16 h photoperiod, *DAM3* expression was found to increase as well, although at a much reduced rate compared to the 10 h photoperiod. Similarly to *DAM3*, under 10 h photoperiod *DAM4* expression peaked at week 7 and decreased afterwards. Under 16 h photoperiod *DAM4* expression remained mostly stable. There were no clear differences between flowering types (Figure 11).

**Figure 11.**
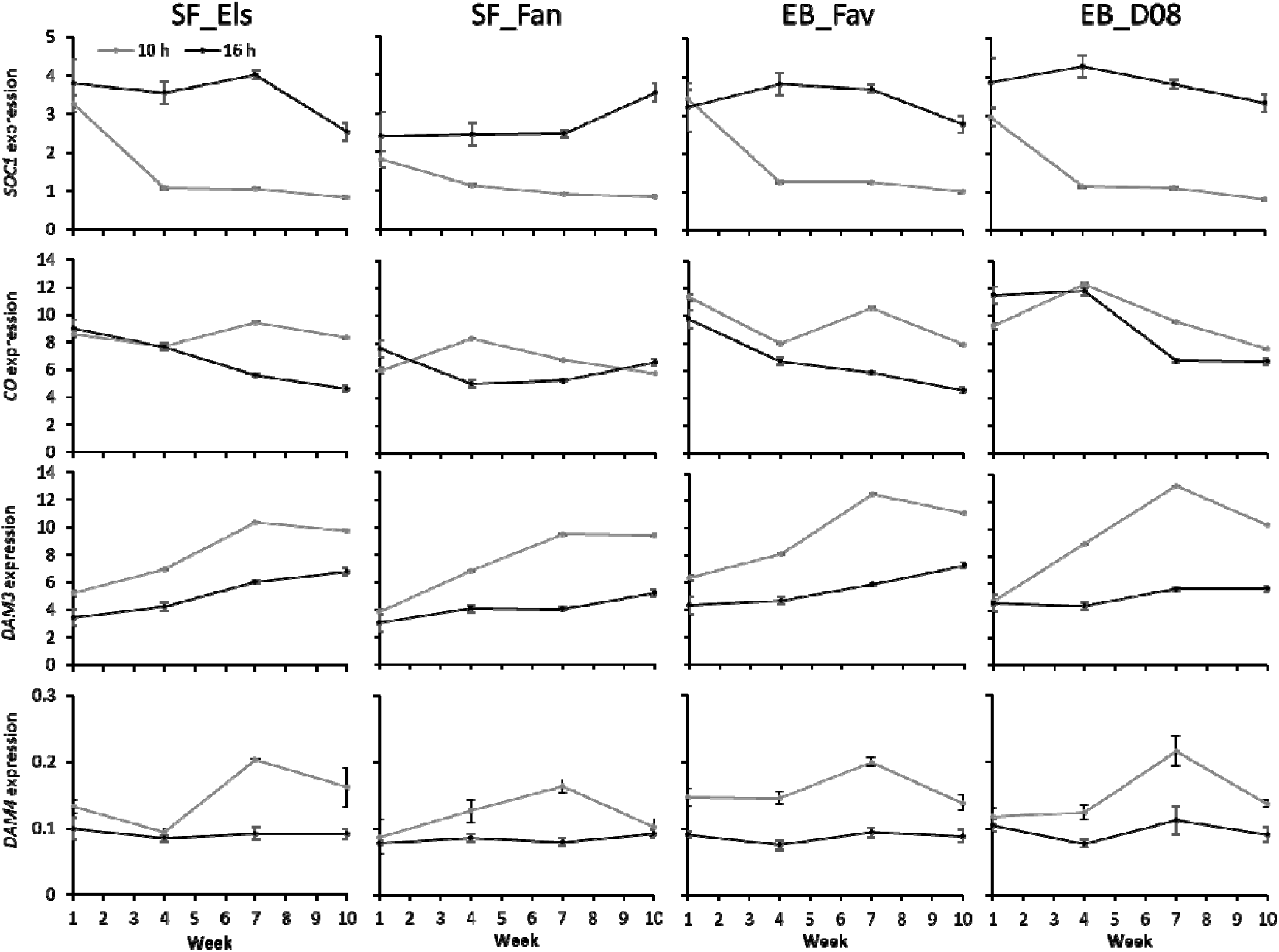
gene expression analysis. On weeks 1, 4, 7, and 10 of the treatment period leaf samples were taken for gene expression analysis. Relative expression levels of the genes *SOC1, CO, DAM3 and DAM4* were determined with qRT-PCR analysis. Expression values were normalized for the expression of the reference genes *DBP*, MSI, and *ACTIN1*. Each data point is the average of three plots, each plot consisting of four replicate plants per cultivar. SF_Els: Elsanta; SF_Fan: Fandango; EB_Fav: Favori; EB_D08: D080-037-2020. Graphs denote the average of three biological replicates ± SEM. Photoperiods of 10 and 12 h represent short day (SD) conditions; 14 and 16 h represent long day (LD) conditions.

## DISCUSSION

The aim of this experiment was to analyze the effect of photoperiod on the early development of seasonal flowering and everbearing strawberry plants. To maintain a constant daily light integral across photoperiod treatments, PPFD was adjusted inversely with day length (Table 2). The responses under investigation, including vegetative morphology, meristem status, and gene expression, are primarily regulated by photoperiod in strawberry (Heide et al. 2013; Stewart and Folta 2010), suggesting that the influence of associated PPFD differences on our interpretation is limited. Nevertheless, at equal DLI, longer photoperiods with lower PPFD can outperform shorter photoperiods with higher PPFD, as photosynthetic efficiency generally decreases with increasing PPFD (Susilo et al. 2025).

### The critical photoperiod for generative development lays between 12 and 14 hours

In seasonal flowering cultivars, endodormancy and flower initiation are both induced under short day conditions (Heide 1977; Sønsteby and Heide 2006, 2021). Under the 10 and 12 h photoperiods, ‘SF_Els’ had decreased petiole length (Figure 3) and leaf area (Figure 4), increased chlorophyll content (Figure 5) and leaf appearance rate (Figure 8), as well as flower initiation (Figure 9). The developmental and physiological responses that we observed are consistent with the induction of endodormancy and indicate a short day developmental response under these photoperiods (Åström et al. 2015; Sønsteby and Heide 2021). In ‘SF_Els’, no generative development occurred under the 14 and 16 h photoperiods, indicating a long day developmental response. Little to no difference was found in vegetative and generative development between the short photoperiods (10 and 12 h) or the long photoperiods (14 and 16 h), contrasting with the results of Heide (1977), who reported a continuous response of petiole length on photoperiod in multiple cultivars (Heide 1977). It should be noted, however, that photoperiodic responses are cultivar specific and parental background can influence outcomes, limiting direct comparison. The destructive harvest after the treatment period yielded a result similar to the weekly measurements. The results indicate a qualitative response of vegetative development to photoperiod with the critical daylength between 12 and 14 h.

### Long day conditions improved vegetative growth in all cultivars

Under the 14 and 16 h photoperiods (long day conditions), we observed increased vegetative growth in the four cultivars included in the experiment. The leaf area of individual leaves was generally smaller in the everbearing cultivars compared to ‘SF_Els’ (Figure 4). However, both everbearing cultivars produced more leaves than ‘SF_Els’, which ultimately resulted in a higher total leaf area (Figure 8). Under long day conditions, petiole length of newly emerged leaves of the everbearing cultivars was typically as high or even higher than that of ‘SF_Els’. Petiole length is considered an important trait, as proper petiole extension may improve light interception (Tan et al. 2022). Moreover, petiole length is often used as a measure of general vegetative growth in strawberry research (Guttridge 1958; Heide 1977; Sønsteby and Heide 2006, 2011).

Photoperiodic effect on vegetative growth is reduced in the everbearing cultivars

Long day conditions improved vegetative growth in ‘EB_Fav’ and ‘EB_D08’, compared to short day conditions. The differences in vegetative growth between short- and long-day conditions were less extreme in these everbearing cultivars. Under short day conditions, the petiole length of the everbearing cultivars increased until the end of the treatment period, albeit at a reduced rate compared to long day conditions (Figure 3). While long day conditions increased the leaf area of individual leaves in the everbearing cultivars, the photoperiodic effect was much smaller compared to ‘SF_Els’ (Figure 4). In contrast to ‘SF_Els’, there was no clear negative effect of long day conditions on the number of leaves produced in the everbearing cultivars (Figure 8). This coincides with the findings of Rivero *et al*. (2021) in ‘EB_Fav’ and a third everbearing cultivar, ‘Delizzimo’ (Rivero et al. 2021). Cultivar-specific differences in photoperiodic responses, including at the level of gene expression, are likely influenced by parental background and should be considered when comparing results across studies. The decreased photoperiod effect on vegetative development in everbearing types resulted in more similar looking plants between short and long day conditions compared to ‘SF_Els’ (Supplement 6). Interestingly, not all vegetative traits had reduced sensitivity to photoperiodic regulation in the everbearing types. Leaf chlorophyll content (Figure 5), runner production (Figure 6), total leaf area and leaf dry weight (Figure 8) all responded similarly to photoperiodic regulation in both flowering types. The everbearing phenotype is the result of a genetic mutation which alters the regulation of generative development (Perrotte et al. 2016; Hytönen and Kurokura 2020). The precise nature of this mutation is as of yet not fully understood. The differences in plant development we observed could be a result of the genetic difference which causes the everbearing flowering type. However, findings from two everbearing cultivars are not necessarily representative of all everbearing cultivars and might be unrelated to the locus which causes everbearing behavior. The physiological basis of this reduced sensitivity, including potential differences in assimilate allocation and source-sink dynamics between flowering types, remains to be investigated.

### Effect of photoperiod on flowering

Photoperiodic treatments did not affect the rate of flowering in both everbearing cultivars (Figure 7). An earlier study found that ‘EB_Fav’ did not flower during 10 weeks of pretreatment at long day conditions (Rivero et al. 2021). This suggests that the treatment period of the current study may have been too short to show the photoperiodic effect, and that the flowers which emerged were initiated during the pre-treatment (which applied the same photoperiod to all subsequent treatment groups). It should be noted that the pre-treatment period was intended for acclimation rather than flower induction. The treatment period itself served as the inductive period, and although ten weeks was insufficient to observe flower emergence in seasonal flowering cultivars, flower initiation was captured through meristem analysis at the end of the experiment (Figure 10). Flowering results in everbearing cultivars should therefore be interpreted as reflecting pre-treatment conditions rather than photoperiodic induction during the experiment. Interestingly, flower emergence was not affected by photoperiod, but runner production was reduced, especially in ‘EB_Fav’. After flower initiation these plants could be grown under short day conditions to prevent runnering. This would affect vegetative growth, but it would also reduce the need for removal of runners, which might be beneficial in circumstances where manual intervention needs to be kept to a minimum.

The seasonal flowering cultivars had no generative meristem at the start of the treatment period, implying that subsequent generative growth had initiated during the treatment period. This indicates that the photoperiod of 13 hours that we applied during acclimation before the treatment period caused a long day response in these cultivars (Figure 9A). A clear difference in flower initiation during the treatment period was found between short day (10 and 12 h) and long day conditions (14 and 16 h) in both seasonal flowering cultivars (Figure 9B-C). No difference in generative growth was found between the two short photoperiods, or between the two long photoperiods. Although ‘SF_Els’ and ‘SF_Fan’ initiated flowers under short day conditions, few to no flowers emerged on these plants, reinforcing the possibility that the treatment period of ten weeks was too short to show an effect on flower emergence, and that conclusions on flowering should be interpreted as early developmental responses rather than full reproductive behavior.

### Gene expression analysis

The expression level and patterns of several genes involved in the regulation of the balance between vegetative and generative growth were analyzed for differences between seasonal flowering and everbearing types.

*SOC1* is a transcription factor which integrates environmental signals and negatively regulates generative growth by inducing expression of the anti-florigen *TFL1*. This gene has been shown to promote vegetative development in wild as well as cultivated strawberry (Mouhu et al. 2013; Muñoz-Avila et al. 2022). *SOC1* expression in the first week of the experimental period was similar between the short day and long day treatments, likely due to the pre-treatment conditions which were equal for all plants (Figure 10). In the short-day treatment, *SOC1* expression reached its lowest point by week four of the treatment period. In ‘SF_Els’ plants grown under short day conditions, this coincided with the peak of leaf area and petiole length (Figure 3, Figure 4). This is consistent with the role of *SOC1* in promoting vegetative development, as reduced *SOC1* expression under short day conditions coincided with the suppression of vegetative growth observed in all cultivars. Contrasting with the other cultivars, in ‘SF_Fan’ an increase in *SOC1* expression was observed by the end of the treatment period, but it is unclear if gene expression has been affected by tip burn, which affected most of the leaves of this cultivar.

Arabidopsis (*At*), *CO* is a key regulator of photoperiodism. Under long days the CO protein induces flowering through the florigen *FT1* (Valverde et al. 2004). Research on woodland strawberry (*F. vesca*) indicates it is likely involved in photoperiodism in strawberry as well (Kurokura et al. 2017; Prisca et al. 2022). However, the expression pattern of strawberry *CO* does not coincide with that of *AtCO*, and the exact role of *CO* in the photoperiodic regulation of development in strawberry is still unclear (Muñoz-Avila et al. 2022). Muñoz-Avila (2022), found significantly higher *CO* expression in long day conditions compared to short day conditions (Muñoz-Avila et al. 2022). Contrastingly, we generally found *CO* to be expressed lower in long day conditions, and in multiple instances no difference was found (Figure 10). A notable difference is that Muñoz-Avila et al., (2022), used the seasonal flowering cultivar Chandler and the perpetual flowering cultivar Selva, which differ from the cultivars used in the present study. Additionally, the plants were exposed to the natural photoperiod. Plants were sampled in December and June to obtain samples from short-day and long-day grown plants. This means that there is a difference in plant age and developmental stage between the short day and long day treatment which may explain the discrepancy between their findings and ours. The absence of a consistent photoperiodic effect on *CO* expression in our data suggests *CO* may not be the primary driver of the developmental differences we observed between short- and long-day conditions.

*DAM3* and *DAM4* are homologs of the *Dormancy-associated MADS-box* genes, first identified in peach (Rodriguez-A et al. 1994; Bielenberg et al. 2008; Hytönen and Kurokura 2020). Previously, we showed that *DAM3* and *DAM4* have similar expression patterns and that these genes are downregulated in chilling conditions (2°C). We consider these genes candidates for master regulators of endodormancy in strawberry (David et al. 2025). Here we show that *DAM3* and *DAM4* are upregulated under the influence of short-day conditions (Figure 10). Both everbearing cultivars experienced higher peaks in *DAM3* expression under short day conditions than the seasonal flowering cultivars. Interestingly, we also observed a gradual increase in *DAM3* expression under long day conditions in all cultivars, most notably in ‘SF_Els’ and ‘Favori’. The gradual increase in *DAM3* expression observed under 16 h photoperiod was unexpected, as David et al. (2025) showed that DAM3 expression increases under short day conditions and decreases upon chilling. The current findings suggest that *DAM3* expression may be regulated by factors other than photoperiod alone, such as developmental stage or temperature, and warrants further investigation. The upregulation of *DAM3* and *DAM4* under short day conditions across all cultivars aligns with the general suppression of vegetative growth observed under these conditions, consistent with their proposed role in regulating seasonal dimorphism in strawberry (David et al., 2025).

### Cultivar-specific vulnerability to tip burn in vertical farming systems environment

During the treatment period, some plants developed leaf tip burn. We responded by increasing nighttime relative humidity (Supplements 4 and 5). Higher RH at night lowers VPD and improves calcium transport to young leaf margins, which helps prevent tip burn (Kroggel and Kubota 2017). This mostly resolved the tip burn problem. However, ‘SF_Fan’ was most gravely affected by tip burn, and, contrary to the other cultivars, did not recover after the climate settings were adjusted. For that reason, the discussion focusses on the other cultivars with respect to vegetative growth and development.

Under longer daylengths, slightly more leaves were affected by tip burn (Supplement 7). Our observations are an example of interaction between environment and genotype and underline the importance of optimization of both of these factors in vertical farm cultivation. In controlled environments, maintaining low VPD during the night promotes guttation and improves calcium supply to emerging leaves, making nighttime humidity management an important tool to prevent tipburn in genetically susceptible cultivars (Kroggel and Kubota 2017).

## AUTHOR CONTRIBUTION STATEMENT

SD: Conceptualization, Data curation, Methodology, Project administration, Validation, Writing – original draft; GF: Methodology, Validation; LFMM: Acquisition, Conceptualization, Supervision, Writing – review and editing; JCV, Conceptualization, Supervision, Writing – review and editing.

## DECLARATION OF INTEREST STATEMENT

The authors report no competing interests related to this work.

## DATA AVAILABILITY STATEMENT (DAS)

The authors confirm that the data supporting the findings of this study are available within the article [and/or] its supplementary materials.

## FUNDING SOURCES

This research is part of the TTW Perspectief programme “Sky High”, which is supported by AMS Institute, Bayer, Bosman van Zaal, Certhon, Fresh Forward, Grodan, Growy, Own Greens/Vitroplus, Priva, Signify, Solynta, Unilever, van Bergen Kolpa Architects, and the Dutch Research Council (NWO).

## SUPPLEMENTS

**Supplement 1.**
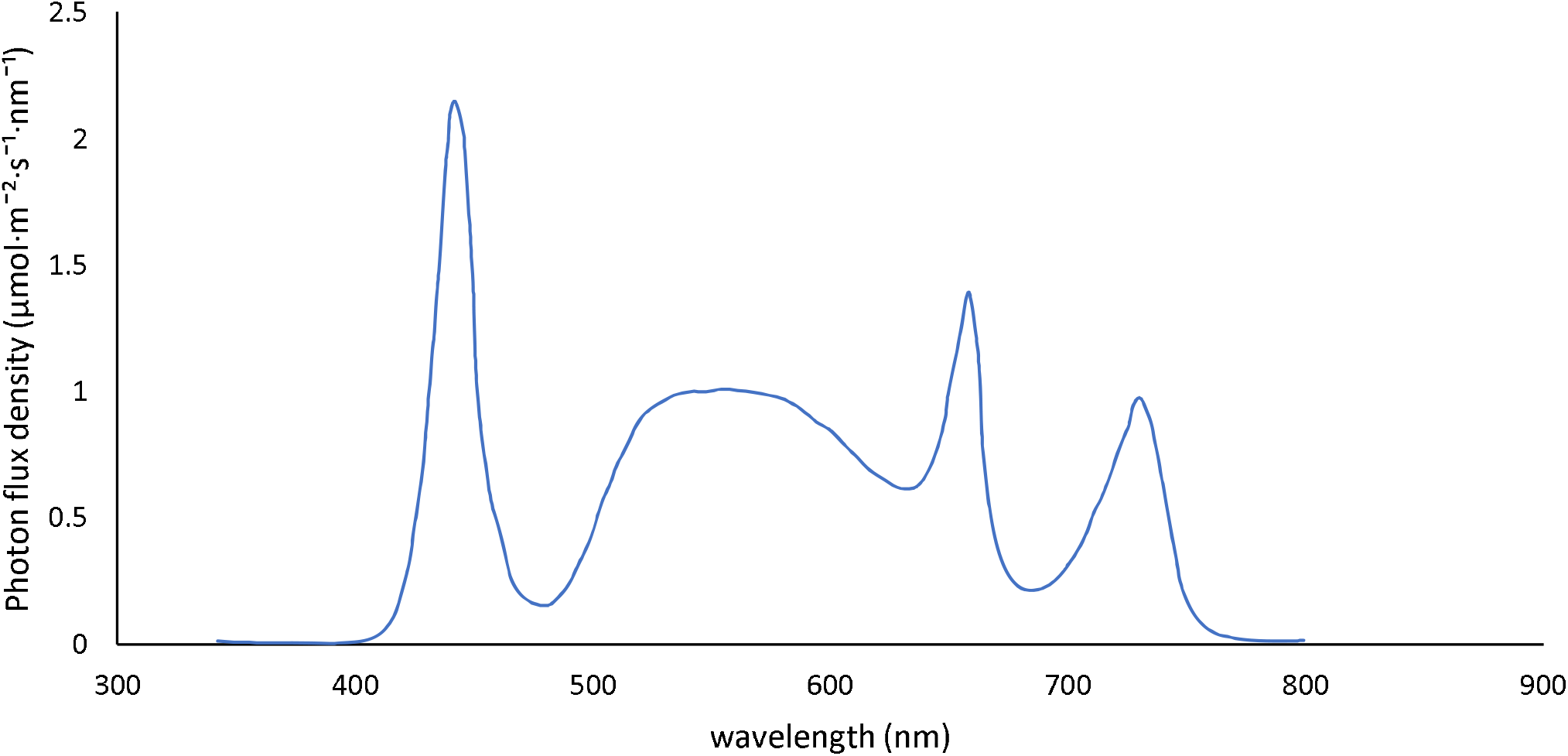
Light spectrum measured with a spectroradiometer.

**Supplement 2.**
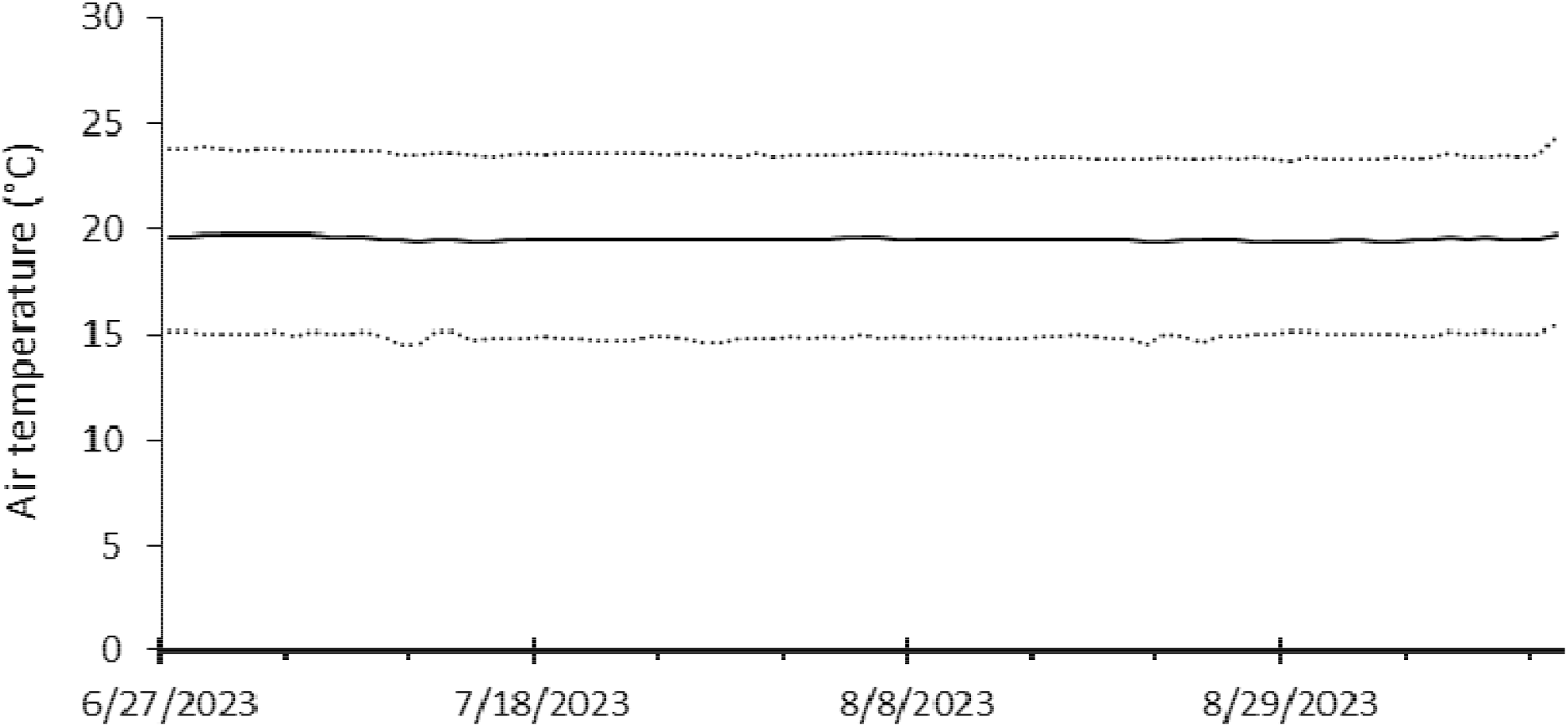
Daily air temperature for the duration of the experiment. Solid line is the average. Dotted lines are the minimum and maximum measured values. Data from each compartment were averaged.

**Supplement 3.**
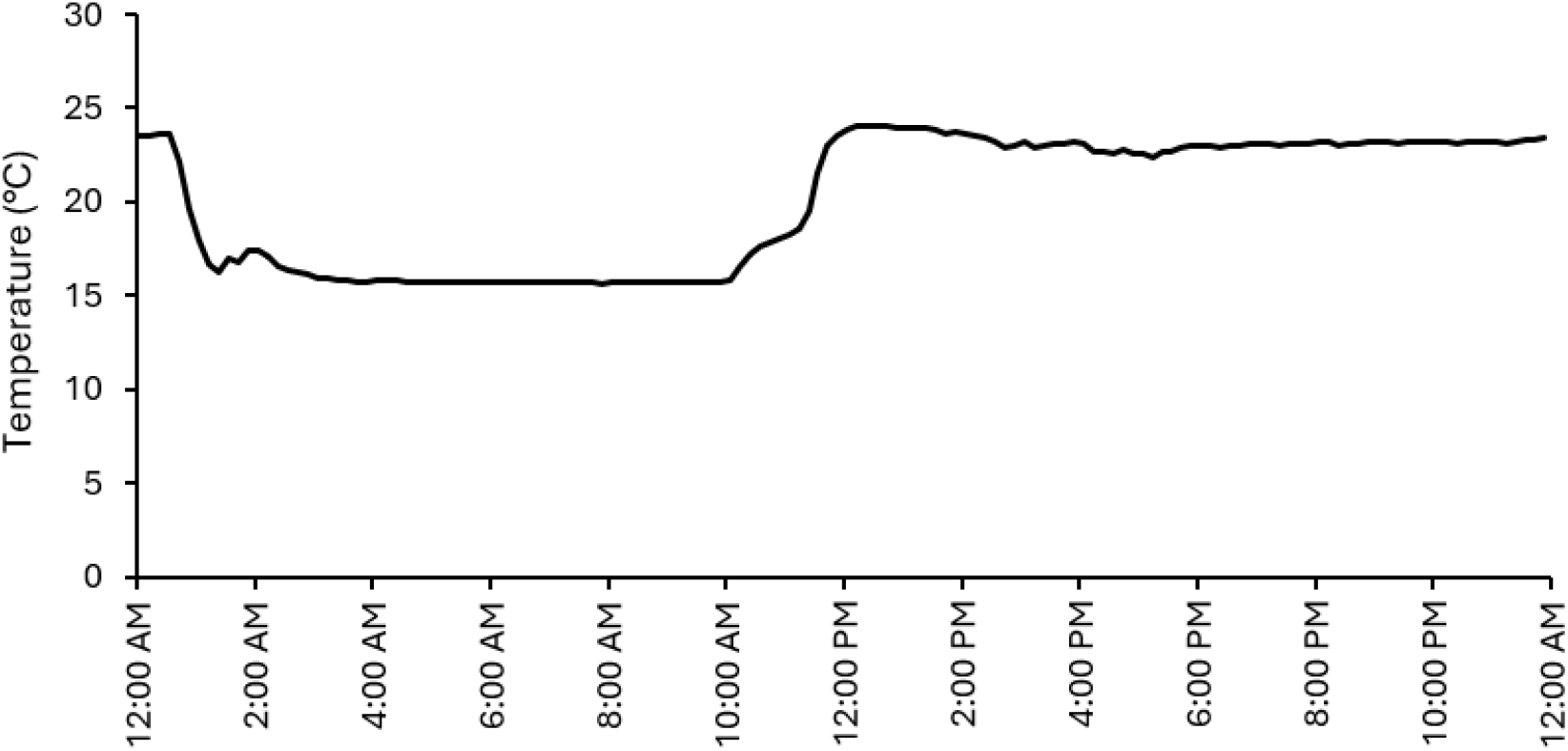
Air temperature per 10 minutes, on July 4, 2023. Data shown is from compartment 5. The dynamic of temperature fluctuation throughout the day is representative of all compartments.

**Supplement 4.**
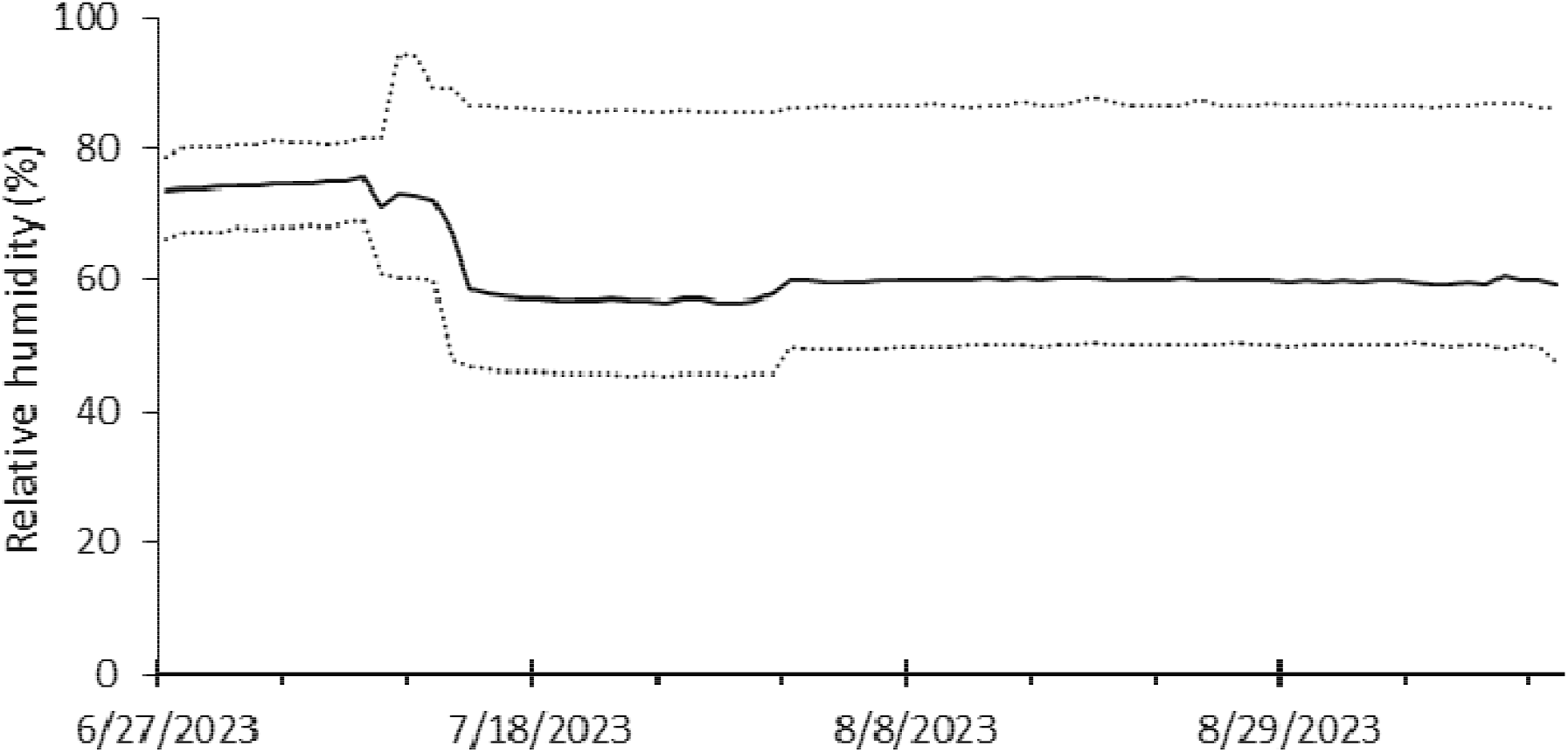
Daily relative humidity for the duration of the experiment. Solid line is the average. Dotted lines are the minimum and maximum measured values. Data from each compartment were averaged.

**Supplement 5.**
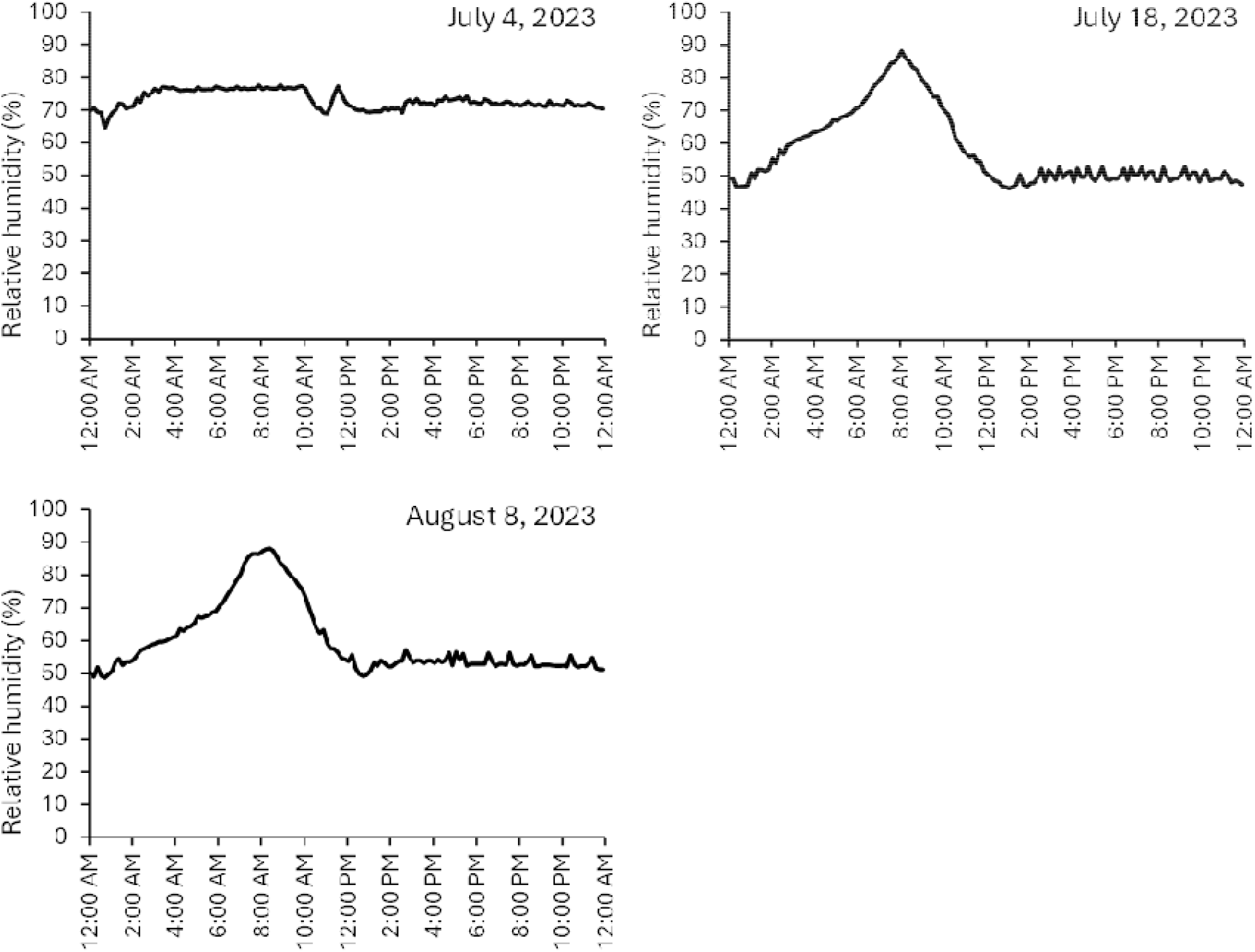
Relative humidity per 10 minutes, on July 4, July 18 and August 8, 2023. Data shown is from compartment 5. The dynamic of relative humidity fluctuation throughout the day is representative of all compartments.

**Supplement 6.**
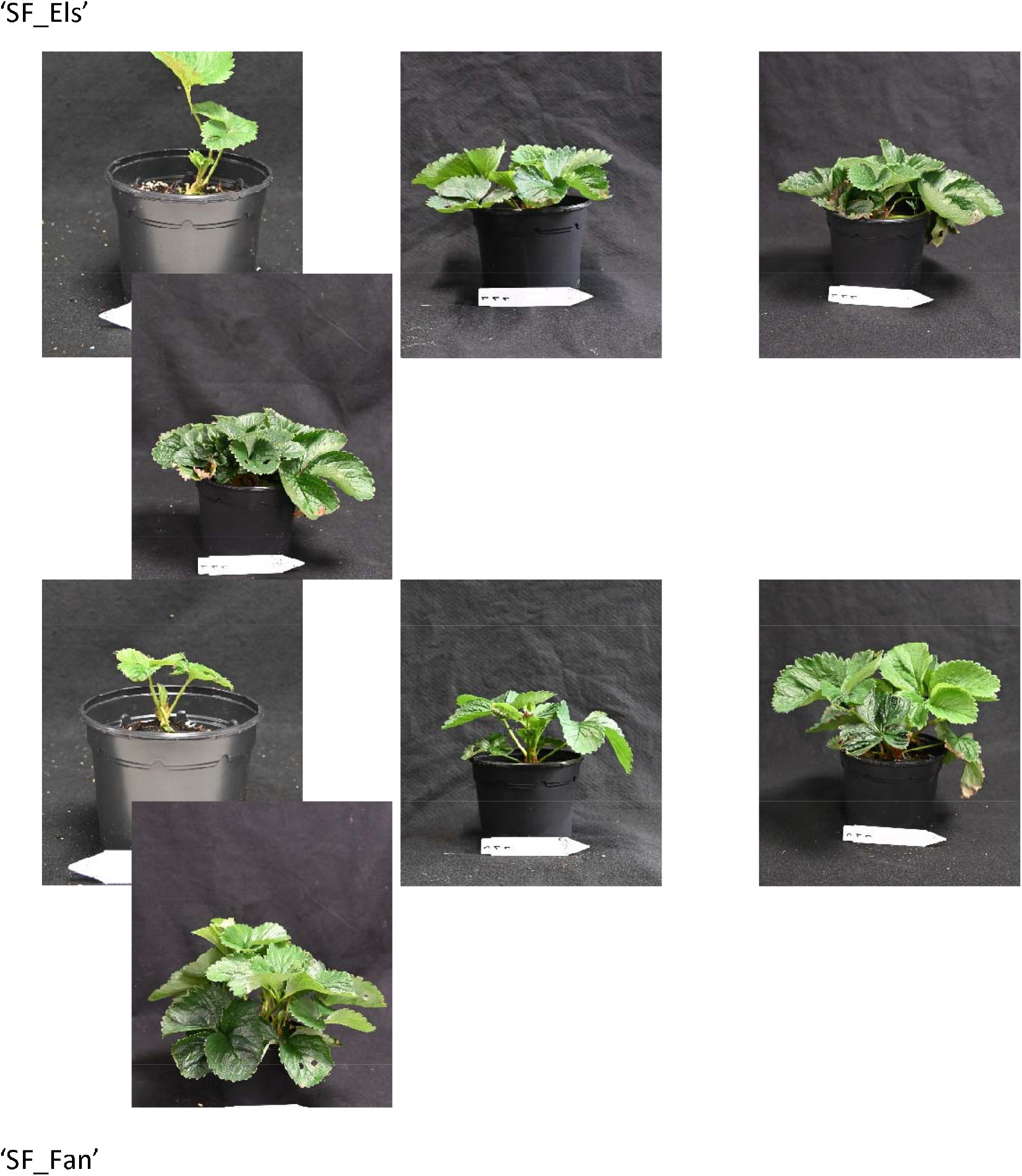
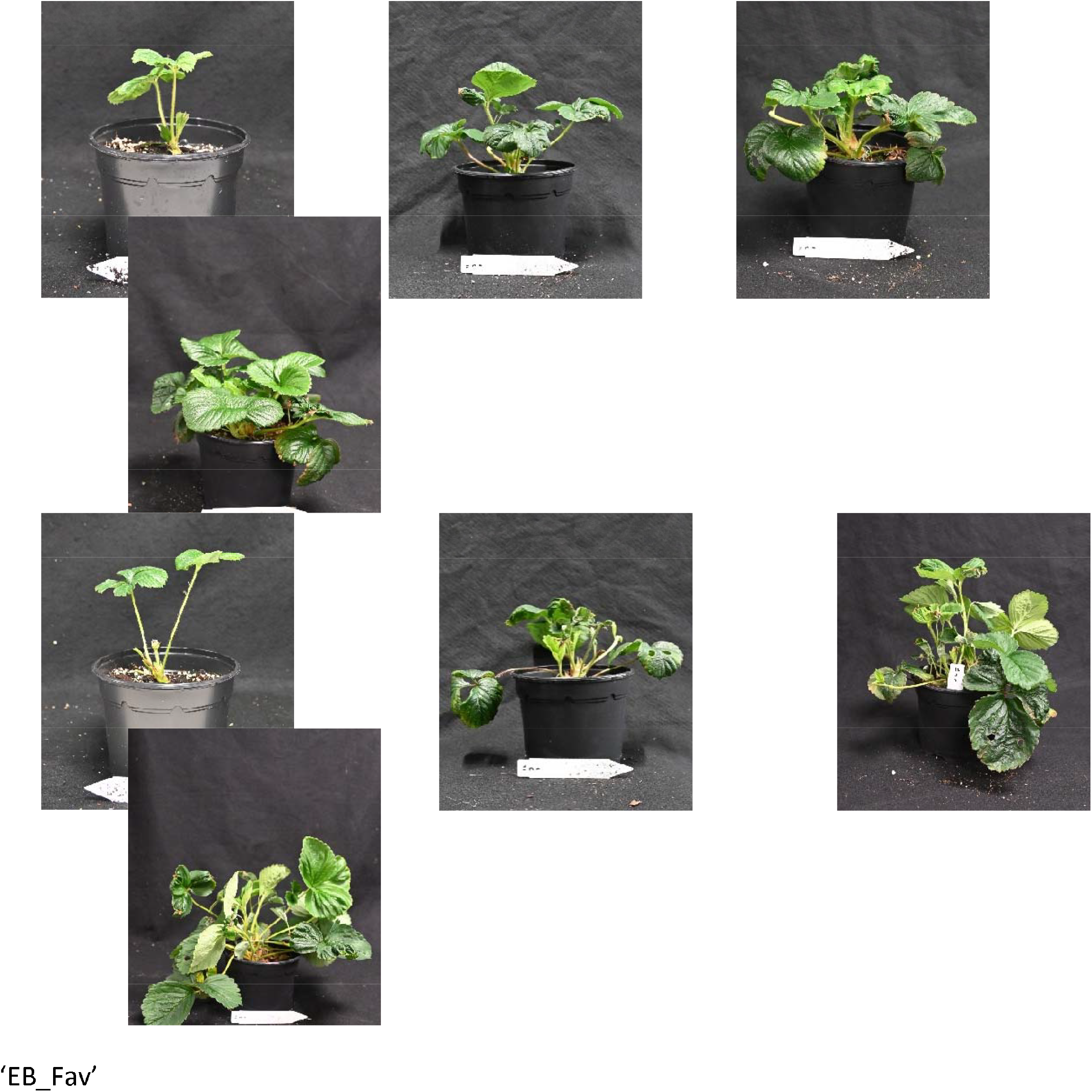
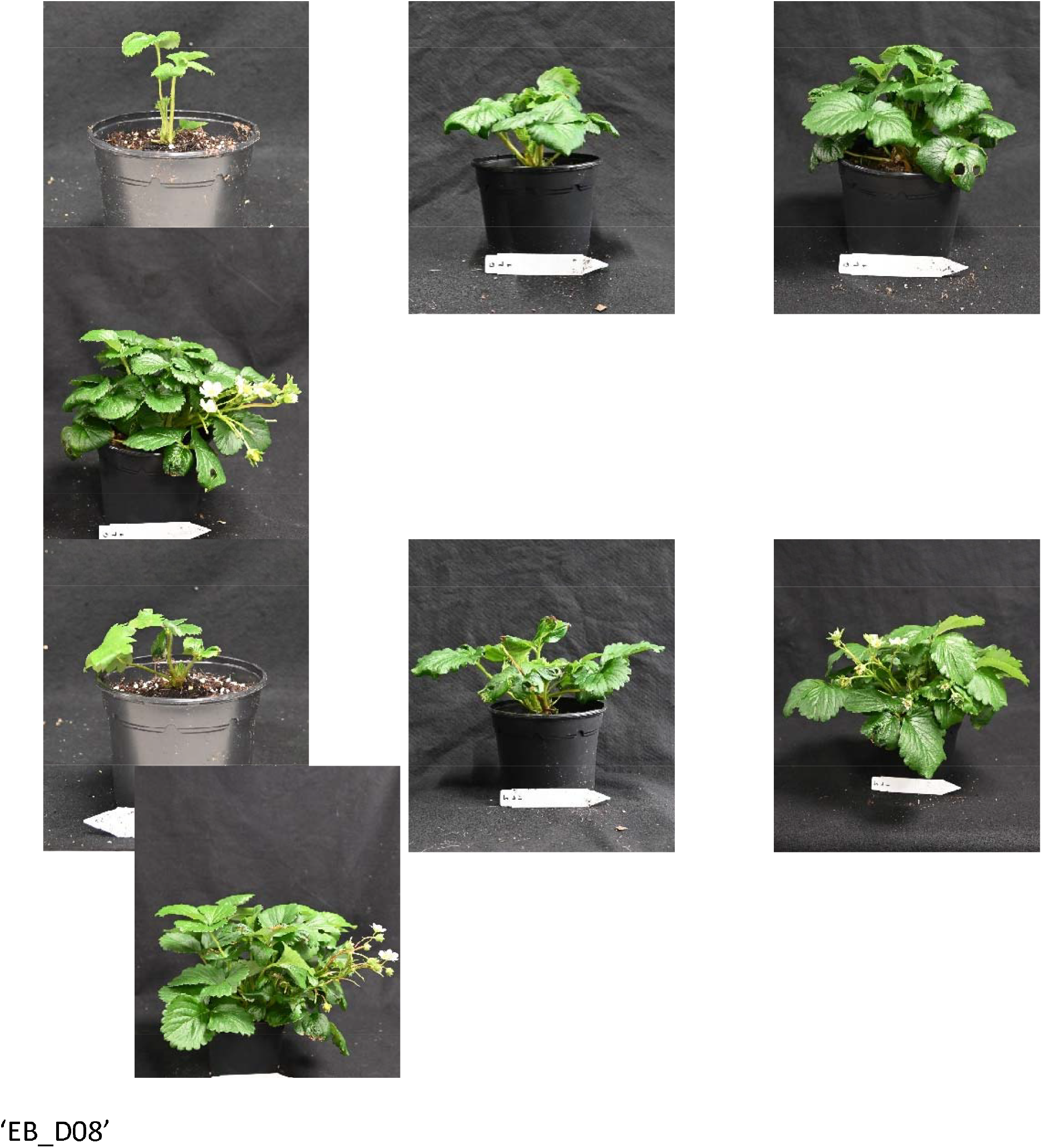
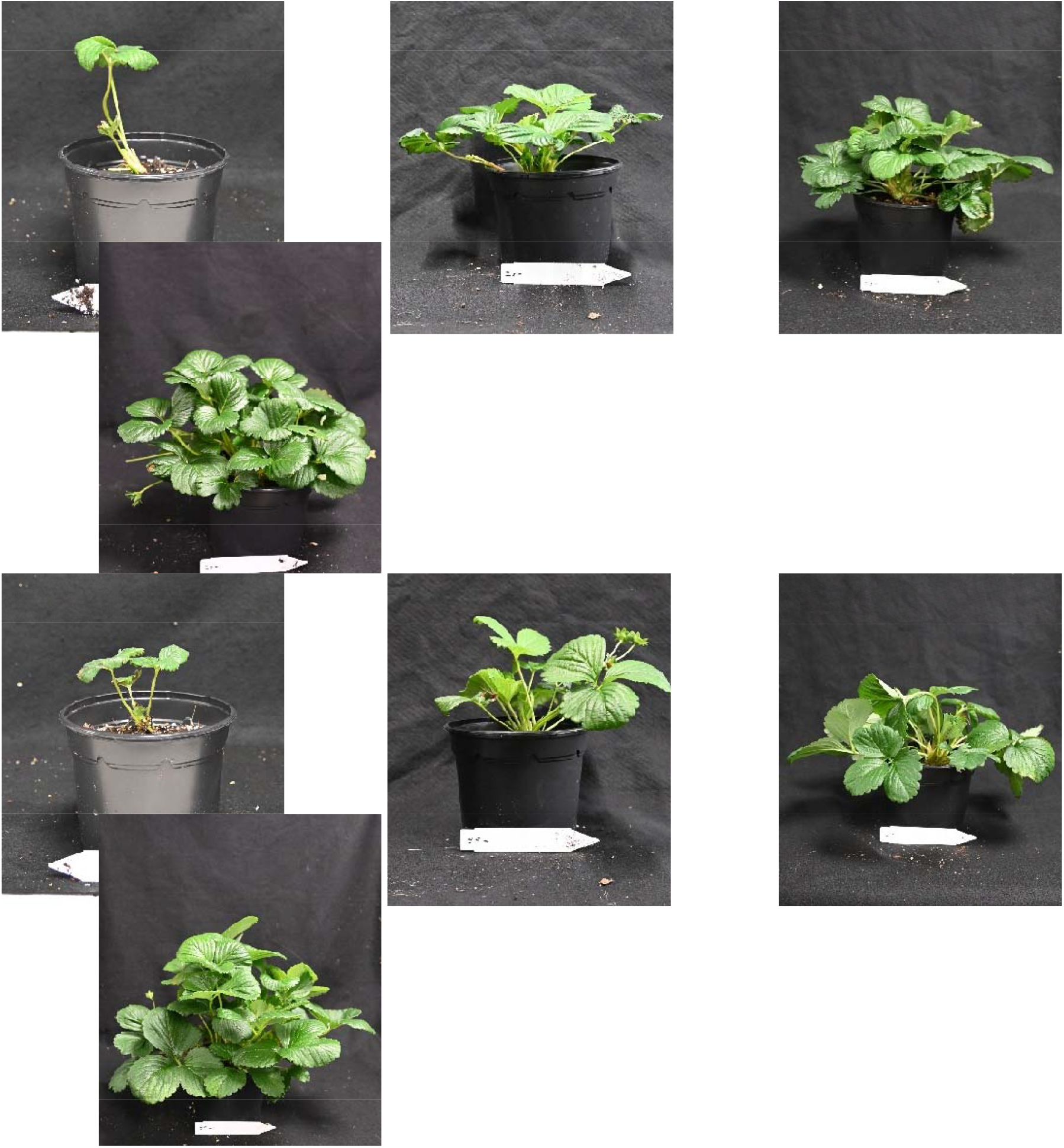
top rows: short day conditions (10 h); bottom rows: long day conditions (16 h). From left to right: week 1; week 4; week 7; week 10. Each time series contains four pictures of the same plant. Appearance of the plants under 10 h and 16 h photoperiod is representative of those under 12 h and 14 h photoperiod, respectively, in terms of morphology.

**Supplement 7.**
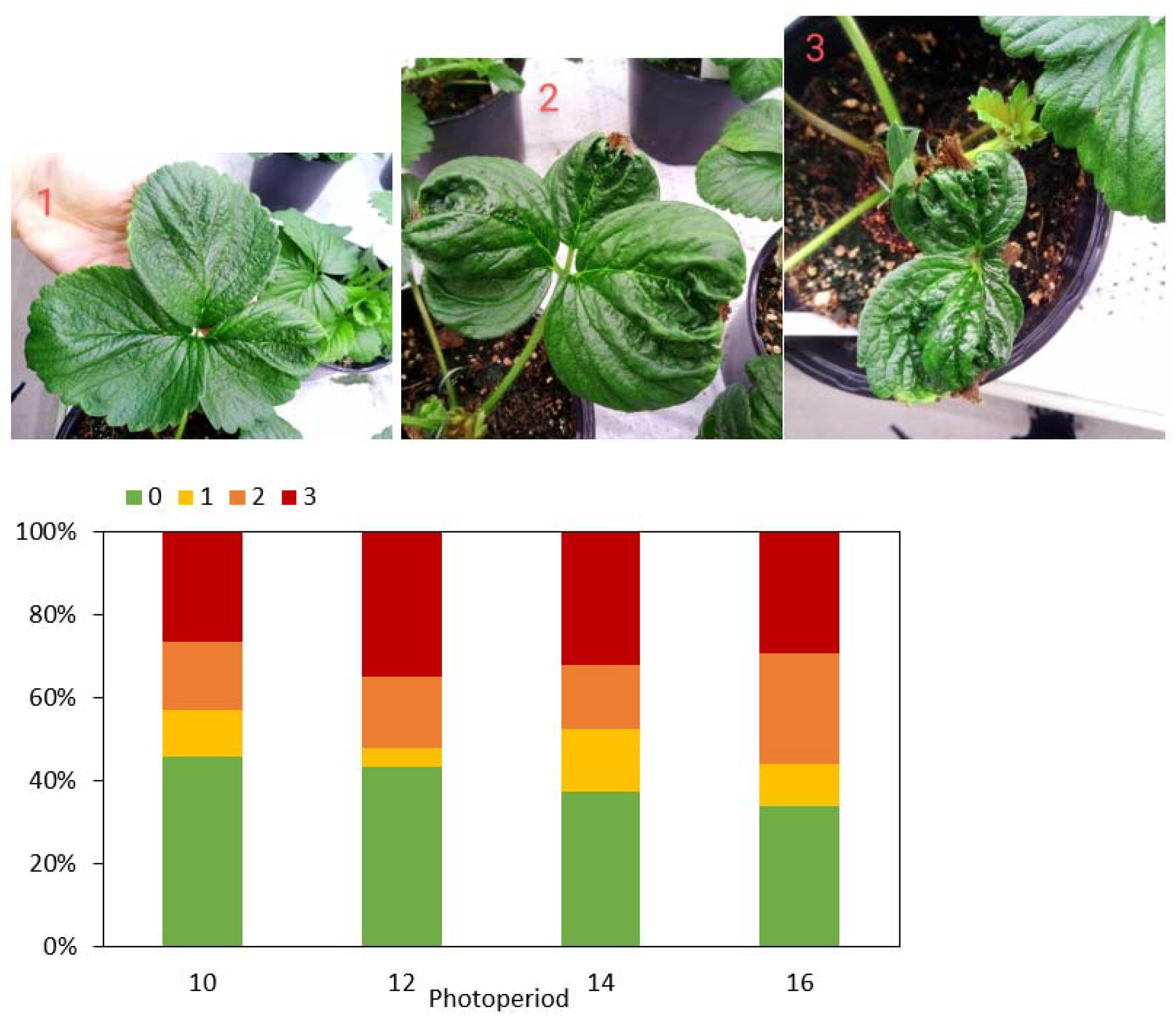
Severity of tip burn in ‘SF_Fan’. The leaves of all plants were assessed on 20/7/2023 and graded on a scale from 0 to 4 (0: no tip burn; 1: slight damage of one or two leaf lobes; 2: more than half of at least one lobe is damaged; 3: the leaf has not expanded due to tip burn). The percentage of the occurrence of each number on the scale was calculated for each photoperiod.

## Notes

### Competing Interest Statement

The authors have declared no competing interest.

### Summary of Updates

fixed author name error due to accent

## REFERENCES

Åström H, Metsovuori E, Saarinen T, Lundell R, and Hänninen H. Morphological characteristics and photosynthetic capacity of Fragaria vesca L. winter and summer leaves. Flora - Morphology, Distribution, Functional Ecology of Plants. 2015:215:33–39. 10.1016/j.flora.2015.07.001

Bielenberg DG, Wang Y (Eileen), Li Z, Zhebentyayeva T, Fan S, Reighard GL, Scorza R, and Abbott AG. Sequencing and annotation of the evergrowing locus in peach [Prunus persica (L.) Batsch] reveals a cluster of six MADS-box transcription factors as candidate genes for regulation of terminal bud formation. Tree Genetics & Genomes. 2008:4 (3):495–507. 10.1007/s11295-007-0126-9

David S, Han J, Marcelis LFM, and Verdonk JC. FaDAM3 and FaDAM4 are candidate genes for the regulation of seasonal dimorphism in cultivated strawberry. 2025:2024.12.10.627498. 10.1101/2024.12.10.627498

De Camacaro MEP, Camacaro GJ, Hadley P, Battey NH, and Carew JG. Pattern of Growth and Development of the Strawberry Cultivars Elsanta, Bolero, and Everest. jashs. 2002:127 (6):901–907. 10.21273/JASHS.127.6.901

CBS (Centraal Bureau voor de Statistiek). Groenteteelt; oogst en teeltoppervlakte per groentesoort. 2024. https://www.cbs.nl/nl-nl/cijfers/detail/37738. Retrieved February 14, 2025.

van Delden SH, SharathKumar M, Butturini M, Graamans LJA, Heuvelink E, Kacira M, Kaiser E, Klamer RS, Klerkx L, Kootstra G, et al. Current status and future challenges in implementing and upscaling vertical farming systems. Nat Food. 2021:2 (12):944–956. 10.1038/s43016-021-00402-w

Demirsoy H, Demirsoy L, and Ozturk A. Improved model for the non-destructive estimation of strawberry leaf area. 101051/fruits:2005014. 2005:60. 10.1051/fruits:2005014

Guttridge CG. The Effects of Winter Chilling on the Subsequent Growth and Development of The Cultivated Strawberry Plant. Journal of Horticultural Science. 1958:33 (2):119–127. 10.1080/00221589.1958.11513920

Heide OM. Photoperiod and Temperature Interactions in Growth and Flowering of Strawberry. Physiologia Plantarum. 1977:40 (1):21–26. 10.1111/j.1399-3054.1977.tb01486.x

Heide OM, Stavang JA, and Sønsteby A. Physiology and genetics of flowering in cultivated and wild strawberries – a review. The Journal of Horticultural Science and Biotechnology. 2013:88(1):1–18. 10.1080/14620316.2013.11512930

Hytönen T and Kurokura T. Control of Flowering and Runnering in Strawberry. Hort J. 2020:89 (2):96–107. 10.2503/hortj.UTD-R011

Kroggel M and Kubota C. Controlled environment strategies for tipburn management in greenhouse strawberry production. Acta Hortic. 2017:(1156):529–536. 10.17660/ActaHortic.2017.1156.78

Kouloumprouka Zacharaki A, Monaghan JM, Bromley JR, and Vickers LH. Opportunities and challenges for strawberry cultivation in urban food production systems. PLANTS, PEOPLE, PLANET. 2024:6 (3):611–621. 10.1002/ppp3.10475

Kurokura T, Samad S, Koskela E, Mouhu K, and Hytönen T. Fragaria vesca CONSTANS controls photoperiodic flowering and vegetative development. Journal of Experimental Botany. 2017:68 (17):4839–4850. 10.1093/jxb/erx301

Lewers KS, Castro P, Hancock JF, Weebadde CK, Die JV, and Rowland LJ. Evidence of epistatic suppression of repeat fruiting in cultivated strawberry. BMC Plant Biol. 2019:19(1):386. 10.1186/s12870-019-1984-7

Maeda K and Ito Y. Effect of Different PPFDs and Photoperiods on Growth and Yield of Everbearing Strawberry ‘Elan’ in Plant Factory with White LED Lighting. Environ Control Biol. 2020:58(4):99–104. 10.2525/ecb.58.99

Mouhu K, Hytönen T, Folta K, Rantanen M, Paulin L, Auvinen P, and Elomaa P. Identification of flowering genes in strawberry, a perennial SD plant. BMC Plant Biol. 2009:9 (1):122. 10.1186/1471-2229-9-122

Mouhu K, Kurokura T, Koskela EA, Albert VA, Elomaa P, and Hytönen T. The Fragaria vesca Homolog of SUPPRESSOR OF OVEREXPRESSION OF CONSTANS1 Represses Flowering and Promotes Vegetative Growth. Plant Cell. 2013:25 (9):3296–3310. 10.1105/tpc.113.115055

Muñoz-Avila JC, Prieto C, Sánchez-Sevilla JF, Amaya I, and Castillejo C. Role of FaSOC1 and FaCO in the seasonal control of reproductive and vegetative development in the perennial crop Fragaria × ananassa. Frontiers in Plant Science. 2022:13.

Perrotte J, Gaston A, Potier A, Petit A, Rothan C, and Denoyes B. Narrowing down the single homoeologous FaPFRU locus controlling flowering in cultivated octoploid strawberry using a selective mapping strategy. Plant Biotechnol J. 2016:14 (11):2176–2189. 10.1111/pbi.12574

Prisca M, Maarten V, Jan VD, Bart N, Wouter S, Timo H, Barbara DC, and Bram V de P. Blue and far-red light control flowering time of woodland strawberry (Fragaria vesca) distinctively via CONSTANS (CO) and FLOWERING LOCUS T1 (FT1) in the background of sunlight mimicking radiation. Environmental and Experimental Botany. 2022:198:104866. 10.1016/j.envexpbot.2022.104866

Rivero R, Remberg SF, Heide OM, and Sønsteby A. Environmental regulation of dormancy, flowering and runnering in two genetically distant everbearing strawberry cultivars. Scientia Horticulturae. 2021:290:110515. 10.1016/j.scienta.2021.110515

Rodriguez-A J, Sherman WB, Scorza R, Wisniewski M, and Okie WR. Évergreen’ Peach Its Inheritance and Dormant Behavior. Journal of the American Society for Horticultural Science. 1994:119 (4):789–792. 10.21273/JASHS.119.4.789

Ruijter JM, Ramakers C, Hoogaars WMH, Karlen Y, Bakker O, van den Hoff MJB, and Moorman AFM. Amplification efficiency: linking baseline and bias in the analysis of quantitative PCR data. Nucleic Acids Res. 2009:37 (6):e45. 10.1093/nar/gkp045

Ruijter JM, Ruiz Villalba A, Hellemans J, Untergasser A, and van den Hoff MJB. Removal of between-run variation in a multi-plate qPCR experiment. Biomolecular Detection and Quantification. 2015:5:10–14. 10.1016/j.bdq.2015.07.001

Schaart J, Salentijn E, and Krens F. Tissue-specific expression of the beta-glucuronidase reporter gene in transgenic strawberry (Fragaria x ananassa) plants. Plant Cell Reports. 2002:21 (4):313–319. 10.1007/s00299-002-0514-4

Schultz DJ, Craig R, Cox-Foster DL, Mumma RO, and Medford JI. RNA isolation from recalcitrant plant tissue. Plant Mol Biol Rep. 1994:12 (4):310–316. 10.1007/BF02669273

Sønsteby A and Heide O. Environmental Regulation of Dormancy and Frost Hardiness in Norwegian Populations of Wood Strawberry (Fragaria vesca L.). The European Journal of Plant Science and Biotechnology. 2011:42–48.

Sønsteby A and Heide OM. Dormancy relations and flowering of the strawberry cultivars Korona and Elsanta as influenced by photoperiod and temperature. Scientia Horticulturae. 2006:110 (1):57–67. 10.1016/j.scienta.2006.06.012

Sønsteby A and Heide OM. Dynamics of dormancy regulation in ‘Sonata’ strawberry and its relation to flowering and runnering. CABI Agric Biosci. 2021:2 (1):4. 10.1186/s43170-021-00026-x

Stewart PJ and Folta KM. A Review of Photoperiodic Flowering Research in Strawberry (Fragaria spp.). Critical Reviews in Plant Sciences. 2010:29(1):1–13. 10.1080/07352680903436259

Susilo KR, Eu A, Besemer B, Heuvelink E, de Vos RCHC, and Marcelis LFM. Extended photoperiod improves growth and nutritional quality of pak choi under constant daily light integral. Front Plant Sci. 2025:16. 10.3389/fpls.2025.1621513

Tan T, Li S, Fan Y, Wang Z, Ali Raza M, Shafiq I, Wang B, Wu X, Yong T, Wang X, et al. Far-red light: A regulator of plant morphology and photosynthetic capacity. The Crop Journal. 2022:10 (2):300–309. 10.1016/j.cj.2021.06.007

Valverde F, Mouradov A, Soppe W, Ravenscroft D, Samach A, and Coupland G. Photoreceptor Regulation of CONSTANS Protein in Photoperiodic Flowering. Science. 2004:303 (5660):1003–1006. 10.1126/science.1091761

Verschoor JA, Otma EC, Qiu YT, van Kruistum G, and Hoek J. Controlled Atmosphere Temperature Treatment: Non-Chemical (Quarantine) Pest Control in Fresh Plant Products. ResearchGate. 2015. 10.17660/ActaHortic.2015.1071.30

Zhang Y, Peng X, Liu Y, Li Y, Luo Y, Wang X, and Tang H. Evaluation of suitable reference genes for qRT-PCR normalization in strawberry (Fragaria × ananassa) under different experimental conditions. BMC Molecular Biology. 2018:19. 10.1186/s12867-018-0109-4

